# Identification of a new gregarine parasite in mass mortality events of freshwater pearl mussels (*Margaritifera margaritifera*) in Sweden

**DOI:** 10.1101/2023.09.26.559503

**Authors:** Anders Alfjorden, Ioana Onut Brännström, Niklas Wengström, Arni Kristmundsson, Mahwash Jamy, David Persson, Fabien Burki

**Affiliations:** Uppsala University, Department of organismal biology, Program in Systematic Biology, Norbyvägen 18D, 752 36 Uppsala, Sweden; National Veterinary Institute (SVA), Department of animal health and antimicrobial strategies, 75189 Uppsala, Sweden; Department of Ecology and Genetics, Evolutionary Biology Center, Uppsala University, 752 36 Uppsala, Sweden; Natural History Museum, University of Oslo, 0562 Oslo, Norway; Swedish Anglers Association, Sjölyckan 6, 416 55 Gothenburg, Sweden; University of Gothenburg, Department of Biological and Environmental Sciences, Box 463, SE-405 30 Gothenburg, Sweden; Institute for Experimental Pathology at Keldur, University of Iceland, Keldnavegur 3, IS-112, Reykjavik, Iceland; Science for Life Laboratory, Uppsala University, Uppsala, Sweden

**Keywords:** *Margaritifera margaritifera*, Freshwater Perl Mussel, Gregarine, Nematopsis, mass mortality, mortality, phylogeny, pathology, ISH, TEM

## Abstract

Freshwater bivalves play key ecological roles in lakes and rivers, largely contributing to healthy ecosystems. The freshwater pearl mussel, *Margaritifera margaritifera,* is found in Europe and on the East coast of North America. Once common in oxygenated streams, *M. margaritifera* is rapidly declining, and consequently assessed as a threatened species worldwide. Deterioration of water quality has been considered the main factors for the mass mortality events affecting this species. Yet, the role of parasitic infections has not been investigated. Here, we report the discovery of three novel protist lineages found in Swedish populations of *M. margaritifera* belonging to one of the terrestrial groups of gregarines (Eugregarinorida, Apicomplexa). These lineages are closely related–but clearly separated– from the tadpole parasite *Nematopsis temporariae*. In one lineage, which is specifically associated with mortality events of *M. margaritifera,* we found using microscopy and *in situ* hybridization cysts in the gills and other organs of diseased individual, containing single vermiform zoites. This represents the first report of a parasitic infection in *M. margaritifera* that may be linked to the decline of this mussel species. We propose a tentative life cycle with distribution of different developmental forms and potential exit from the host into the environment.

## INTRODUCTION

A large diversity of protistan parasites is known to affect invertebrate hosts. In marine environments, it is well established that mortalities in bivalve populations (e.g., mussels, oysters, clams, cockles and scallops), either in wild populations or in aquaculture, are often caused by different groups of parasites belonging to Apicomplexa, Perkinsea, or Ascetosporea (Bower, Hervio, and Meyer 1997; Carnegie et al. 2003; Carrasco, Green, and Itoh 2015; Kristmundsson and Freeman 2018; Čižmek et al. 2020). In comparison, much less is known from freshwater systems, even though populations of different mussel species have been declining (Haag 2019; Richard et al. 2020; Aldridge et al. 2023). The decline among different species of freshwater mussels has mainly been explained by a degradation of water quality caused by anthropogenic activities such as dredging, sedimentation, run-offs, eutrophication, and coal mining (Haag 2019; Richard et al. 2020). However, the possibility of infectious agents, such as protistan parasites, has generally not been considered.

The freshwater pearl mussel *Margaritifera margaritifera* inhabits rivers and creeks throughout the northern Holarctic region, which includes population from North America (common name: eastern pearlshell) as well as northern Europe. While other freshwater mussel species have a wide tolerance for different aquatic environments, *M. margaritifera* only lives in highly oxygenated streams with clear running water, which makes it particularly vulnerable to fluctuations of environmental conditions (Henrikson and Söderberg 2020; von Proschwitz and Wengström 2021). Indeed, this species is assessed as globally threatened according to the International Union for Conservation of Nature (IUCN) Red List (Moorkens et al. 2017), while in Sweden it is highly endangered (Westling et al. 2020). Currently, only one third of the Swedish populations are considered healthy with ongoing recruitment, i.e. reproducing populations (Henrikson and Söderberg 2020; von Proschwitz and Wengström 2021).

Swedish populations of *M. margaritifera* have been closely monitored due to their critical status, with an overview of mortality events including some microscopic as well as macroscopic descriptions performed in 2016 and 2017 (Wengström et. al., 2019). From populations experiencing mass mortality events (MME), patterns of pathology in gonads, foot, digestive gland and mantel are described. However, no clear conclusion on the etiology or possible causes for the observations and histopathological changes could be drawn (Wengström et al. 2019). In a new area with MME detected in 2017 (the locality Teåkersälven), we performed initial histopathology on a few moribund mussels which revealed the presence of cysts with similarities to gregarines (Apicomplexa).

Gregarines are a group of parasites infecting almost exclusively invertebrate hosts. They are highly diversified, both morphologically and genetically (Cavalier-smith 2014; Clopton 2002; Leander 2008). However, there is no published report of gregarines in *M. margarifera* nor in any other freshwater mussels and more generally, to our knowledge, no published data are available on apicomplexan parasites in freshwater molluscs. In contrast, several studies have shown that gregarines are common parasites in marine bivalves (Silva et al. 2019; Carlos Azevedo and Padovan 2004; Vivares 1969; Kinne and Lauckner 1983; Hatt 1931; Tuntiwaranuruk et al. 2004; Berrilli et al. 2000; Belafastova 1996). Gregarines from arthropods and annelids are probably also common in freshwater systems, where they are known to infect a variety of insects that live part of there life in aquatic environments (Clopton 2002; Clopton, Cook, and Cook 2007; Clopton 2009; Lantova and Volf 2014; Smith and Cook 2012; Smith-Herron 2015; Tseng 2007). Tadpoles have also been shown to host a gregarine species, which was described as *Nematopsis temporariae* belonging to Eugregarinorida (terrestrial clade I) based on microscopy, ultrastructure, and 18S rDNA phylogenetic analysis (Chambouvet et al. 2016).

To investigate the gregarine-like parasite observed in *M. margaritifera*, we collected specimens at several time points over three years in Teåkersälven experiencing MME, as well as in a reference healthy population (from the locality Stommebäcken). DNA sequencing of the small subunit of the ribosomal DNA (18S rDNA) revealed a novel diversity of gregarines related to the tadpole parasite *N. temporariae* (Chambouvet et al. 2016). Histology, *in situ* hybridization (ISH), and transmission electron microscopy (TEM) on specimens collected from Teåkersälven provide a first report of a parasitic infection, including a putative lifecycle, in the endangered *M. margaritifera*, and more generally the first report of gregarines in freshwater molluscs.

## MATERIALS AND METHODS

### Sample collection

Live specimens of *M. margaritifera* were collected in two close rivers from Sweden: Stommebäcken (N 58.7512057, E 12.2262805) and Teåkersälven upstream, purification site (PS): N 58.7572073, E 12.2311112 and downstream, PS: N 58.7572073, E 12.2311112). Teåkersälven has been experiencing ongoing mortalities (MME) and a rapid decline since 2016. Data summarizing population sizes of both streams are listed in Supplementary Table S1, clearly showing this decline between 2016-2019. The animals collected in Teåkersälven were showing signs of poor health (i.e., laying on top of the substrate, gaping and a slow response to tactile stimuli (Richard et al., 2020)). Sampling from the MME population was performed in September 2018 (5 specimens collected), October 2019 (10 specimens collected) and June 2020 (20 specimens collected), while the healthy reference population at Stommebäcken (no signs of mortalities were observed) was sampled in September 2018 (3 specimens collected), and June 2020 (10 specimens collected). A bathyscope was used to identify the mussel individuals. A permission for collecting limited numbers of freshwater pearl mussels was granted for this study by the regional authority (länssstyrelsen Västra Götaland: 623-28113-2018; 623-37022-2019; 621-22570-2020). The mussels collected in 2018 and 2020 were transported in containers with water from the sampling sites to the National Veterinary Institute (SVA, Uppsala, Sweden), kept cold during transport using cooling blocks. The mussels collected in 2019 were directly preserved at sampling site using 95% ethanol. In the lab, these animals were dissected in 2-3 mm thin cross sections covering the digestive gland region, cut longitudinally in two halves and stored in 95% ethanol for subsequent DNA extraction.

### Macroscopic observations

Macroscopic investigations were made as described in Wengström et al. (2019) and observations are listed in Supplementary Table S2. Specimens collected in 2018 and 2020 were screened for uncharacteristic gross morphological anatomy and biological measures were taken (Supplementary Table S2), while animals from 2019 were only investigated by molecular methods. Before fixation, released fluid and cells from sectioned organs were collected for cytology by tissue imprints and PCR by cotton swabs. The swabs were directly frozen and stored at −20 ° C until DNA extraction.

### Histology, cytology and transmission electron microscopy

Two parallel cross-sections were collected in tandem cuttings to include both digestive gland and gonads from each animal from the MME population (2018, n=5; 2020, n=20) and fixed in Davidssons freshwater fixative for 48 hours. Fixed sections were further processed according to routine histological protocols (Howard et al. 1983) and stained by hematoxylin and eosin (HE). Cytological imprints were stored dry and dark until further processing and stained using Hemocolor rapid stain of blood smears (product number 1.11956, Merck KGaA, Darmstadt, Germany).

Stained sections of each sampled specimen were inspected by microscopy using both low power (Olympus SZ binocular stereo zoom microscope; Tokyo, Japan) and high-power magnification (Nikon labophot; Tokyo, Japan). Photos were taken with a Canon EOS 500D attached to a microscope by a lens-adaptor (Martin Microscope Company MM-SLR adaptor S/N:0468; Easley, SC USA). Additional photos were taken for illustration of the putative live cycle, using an inverted microscope Nikon eclipse Ts2R using Nikon Plan Fluor DIC, 100X/1.30 oil lens and TIS Camera USB 3.0 Monochrome 1/1.2 CMOS. Organ biopsies for TEM were collected from digestive gland and fixed in 2.5% glutaraldehyde solution (0.1 M Phosphate Buffer, pH 7.4 (PB)). Selected samples (2018, n=2; 2020, n=2) were further processed by PB rinsing before and after 1 hour incubation in 1% osmium tetroxide and then dehydrated in graded alcohols (70%-99.9%), followed by incubation in propylene oxide for 5 min. Embedding followed routine protocol, using Epon Resin and propylene oxide (1:1) for 1 hour, two changes of 100% resin and subsequently polymerization in Epon Resin for 2 days. Semithin sections (1-2 microns) were stained by toluidine blue and examined for areas of interest. The block was trimmed and ultrathin sections (60-70 nm) were cut in a Leica UC7 Ultramicrotome and placed on a grid. Sections were contrasted using 5 % uranyl acetate for 10 minutes and 3 % Reynolds lead citrate for 2 minutes. Grids were examined by TEM (FEI Tecnai G2) operated at 80kV.

### DNA extraction and PCR amplification

For the ethanol-preserved animal specimens, the ethanol was evaporated for 10 minutes and rinsed for 30 minutes in 1x PBS solution before the DNA isolation procedure. To break the cell wall of encysted cells and increase the DNA yield, all samples collected from digestive glands were pretreated using freeze – thawing cycles (3X -10 min deep freezing at−70 °C followed by 1 min heat block treatment at 90 °C) and digested with Proteinase K and ATL buffer at 56°C overnight under slow shaking condition (60 rpm). DNA was isolated with Qiagen DNeasy Blood and tissue kit, following manufacturer’s instructions. DNA concentration was measured on a Qubit Fluorometric Quantification Machine (Thermo Fisher Scientific).

The 18S rDNA sequences were obtained in two steps. First, DNA was amplified from all mussels collected in 2018 (n=8) using a general non-metazoan reverse primer 18s-Euk 1134-R (5’-TTTAAGTTTCAGCCTTGC-3’) (Bower et al., 2004) combined with the primer Int F1 (5’-GATTAAGCCATGCATGTCTAAG-3’) known to amplify marine gregarines (Wakeman & Leander, 2013). The PCR reaction was conducted using EconoTaq PLUS 2X Master mix (Cat no. 30035-1, Lucigen, LGC Biosearch technologies, USA) as follow: Initiation, 94 oC for 2 min; 35 cycles of 94 oC for 30 sec, 52 oC for 1 min, 72 oC for 1.5 min; final extension, 72 oC for 10 min. PCR products were cloned using StrataClone PCR cloning kit (product no: 240205, Agilent technologies Sweden AB, Kista) and re-amplified with M13F/M13R primers using DreamtTaq (Thermo Fisher Scientific, Baltics UAB, Vilnius, Lithuania). Bands of the expected size were purified with IllustraTM ExoProStarTM 1 step Enzymatic PCR and Sequence reaction clean up Kit (prod no. US77702, GE Health Care UK limited, Buckinghamshire). Purified clones were Sanger-sequenced in both direction at Macrogen (Macrogen Europe BV, Amsterdam, Netherland). The sequences were verified, trimmed, and assembled with Geneious v.9 (Kearse et al. 2012). Based on these initial sequences, new specific primers were designed to further explore the Nematopsis lineage diversity, on mussels sampled (n=48) between 2018-2020. Primers were designed to cover the v4 region but also to reach almost full length of 18S rRNA using both available NCBI sequences from tadpoles (Chambouvet et al., 2016) and with the initial sequences generated in this study (above). All primers used in our study are listed in Supplementary Table S3. All gregarine 18S rDNA sequences obtained in this study have been deposited in GenBank under the accession numbers (OR133332 - OR133340, OR167014-OR167032, OR167182-OR167194, OR167380-OR167381, OR168121-OR168122, OR168627). The accession numbers of the sequences listed in Supplementary Table S4.

### Environmental diversity

To investigate the environmental phylogenetic diversity of *Nematopsis*, we BLASTed (Altschul et al., 1990) the Sanger sequences against a long-read environmental dataset previously published (Jamy et al. 2022), using a 90% similarity cut-off. Briefly, this dataset was generated by PacBio sequencing of ∼4500 bp of the ribosomal operon (18S, ITS1, 5.8S, ITS2, 28S) of the eukaryotic community from various habitats, including four freshwater samples. Three samples were obtained from Swedish lakes (Lake Erken, Lake Ersjön, and Lake Stortjärn) (Jamy et al. 2022) while one sample was collected from permafrost thaw ponds from Canada (Peura et al., 2020) (Supplementary Table S5). We also analyzed short-read environmental data to investigate the distribution of *Nematopsis* in various habitats using datasets corresponding to Tara Oceans (de Vargas et al., 2015; V9), VAMPs (Huse et al., 2014; V9), Scandinavian Lakes (Khomich, Kauserud, Logares, Rasconi, & Andersen, 2017; V4), Neotropical soils (Mahé et al., 2017; V4), Global soils (Bates et al., 201; V4), Swiss alpine soils (Seppey et al., 2020; V4 and Singer et al., 2020; V9), Lake Baikal (Annenkova, Giner, & Logares, 2020; V4), and St. Charles River (Cruaud et al., 2019; V4). All datasets were cleaned and Amplicon Sequence Variants (ASVs) were generated using DADA2 (Callahan et al., 2016). Sanger sequences and positive hits from the long-read dataset were used to query the ASVs with BLAST (85% similarity threshold), using the V4 and V9 fragments as appropriate (Altschul et al., 1990).

### Phylogenetic analysis

The 18S rDNA sequences obtained from MME and reference populations of *M. margaritifera* were blasted online against the NCBI nucleotide database (last access: May 2023) and representative hits across Apicomplexan diversity, plus publicly available environmental sequences were retrieved. Additionally, reference sequences from all the major Apicomplexan lineages and Chrompodellida were included in the dataset together with Perkinsozoa and Dinophycaea, chosen as outgroup. The OTUs assembled from the long-read environmental sequences described in (Jamy et al. 2022) were also added to the final dataset (see section above). A list of all sequences used for phylogenetic analyses is provided in Supplementary table S6. The sequences were aligned with MAFFT v.7.407 (Rozewicki, Li, Amada, Standley, & Katoh, 2019) using the mafft-linsi algorithm. TrimAl v. 1.4.1 (Capella-Gutiérrez, Silla-Martínez, & Gabaldón, 2009) with gappyout setting was used to trim ambiguously aligned sites. IQTREE v.2.0 -rc2-omp-mpi (Minh et al., 2020) was used to infer a Maximum Likelihood (ML) tree, using the best-fitting model of nucleotide substitutions GTR+F+I+G4 as determined by the corrected akaike information criterion. The branch support was assessed with standard nonparametric bootstrap based on 100 bootstrap replicates. The raw, aligned and trimmed fasta files used for the phylogenetic analysis are available in FigShare (under embargo).

### In-situ hybridization (ISH)

We followed the method described in (Kristmundsson & Freeman, 2018), with one modification using an extra blocking step against endogenous biotin signal. Two biotin labelled probes: 360rev(ISH) 5’-TGGACTGTTGCCAGTCCTTC-Bio and 1143rev(ISH) 5’-TAGACGTATGATTGACGTGC-Bio where designed to specificly target the new gregarine sequences amplified from the MME population of *M. margaritifera* (Teåkersälven) using sequences originating from mussels collected in 2018, i.e., lineage A. Cross-sections from embedded mussels in paraffin blocks were cut at 7 μm and 5 μm thick then deparaffinized, rehydrated in distilled H2O followed by PBS incubation and permeabilized for 12 minutes at 37 °C using 7 μg/ml proteinase K in Tris-buffered saline (TBS) pH 8 followed by washing in PBS. The sections were then incubated in 0.4 % paraformaldehyde in PBS for 15 minutes followed by washing with distilled water. To prevent unspecific binding, we used a streptavidin-biotin blocking kit (Vector Laboratories Inc., Burlingame, USA) by first incubating with streptavidin for 15 minutes, washing in buffer followed by 15 minutes incubation with biotin and washing. This was followed by a secondary blocking of hydrogen peroxide, where 10 % hydrogen-methanol was added for 10 minutes followed by a 5 minutes washing step repeated one time. The hybridization buffer was added with probes using Frame-SealTM enclosure (Bio-Rad, Sundbyberg, Sweden). By adding 100-120 μl of ready-to-use hybridization buffer (Roche prod Merck KGaA, Darmstadt, Germany) mixed with both probes (2.25 ng/ml of each), the sections were sealed, denatured at 95 °C during 4 minutes and hybridized at 45 °C for 8 hours (overnight) in a humid chamber. Negative control slides were incubated in buffer without adding the probes. The hybridization buffer was washed off in two steps with non-stringent and stringent buffer, respectively: first with 2x SSC kept at room temperature followed by second buffer SSC with 0.1 % Tween 20 stored at 45 °C (saline-sodium citrate buffer, SSC). For signal detection the samples were incubated with horseradish peroxidase-labelled streptavidin (Dako, Agilent Technologies, Glostrup, Denmark) for 20 minutes at room temperature followed by 3 × 5 minutes washing in PBS (pH 7.4) and then adding Nova Red Peroxide substrate for 10 minutes (Vector laboratories Inc., Burlingame, USA, Maravai Life Sciences). Hematoxylin was used as counterstain followed by dehydration and mounting of coverglas.

## RESULTS

### Phylogenetic analysis reveals three novel terrestrial gregarine lineages isolated from *M. margaritifera*

After a mass mortality event (MME) of *M. margaritifera* in Teåkersälven first detected in 2017, a general decline of the population was continuously observed (Wengström et al., 2019), culminating to about 65% in 2019 (Supplementary Table S1). To determine the potential role of a parasitic infection in this decline, specimens were collected between 2018 and 2020 from both the MME population (35 specimens collected) and a healthy reference population in a nearby locality (Stommebäcken, 13 specimen collected). A total of 46 sequences were obtained using 18S rDNA primers known to amplify gregarine diversity (see Material and Methods and Supplementary Table S4). To investigate the phylogenetic origin of these sequences, a diverse set of apicomplexan homologs were retrieved by BLAST against the NCBI database, as well as six long-read environmental sequences obtain from (Jamy et al. 2022) (Supplementary Table S5). Only one short-read metabarcoding sequence was identified by our search against various datasets corresponding to a rare OTU from a Norwegian lake (Khomich et al. 2017); because this short sequence was identical to two of the long reads from Swedish lakes, we did not include it in the phylogenetic reconstruction.

Maximum Likelihood (ML) phylogeny showed that all new sequences isolated from *M. margaritifera* are part of the terrestrial clade 1 of gregarines, together with neogregarines, monocystids, and species from insects especially those associated with water (e.g. damselfly, dragonfly, water scorpion) and other environmental sequences (Figure 1, Supplementary Figure S1). The newly generated sequences form three distinct lineages, here named lineages A-C (Figure 1). Lineage A is supported by 99% bootstrap and includes the majority of sequences isolated from the MME population (35/36), one sequence isolated from the reference populations and five long-read environmental sequences from permafrost soil from Canada and Swedish lakes and bogs (Figure 1, Supplementary Figure S1). Lineage A branches together (bootstrap support = 100%) with a clade comprising the previously published sequences of the tadpole parasite *Nematopsis temporariae* (Chambouvet et al. 2016) and an environmental sequence from the Canadian permafrost soil, but also with a distinct sequence (lineage B) isolated from the 2019 MME population in Teåkersälven (Figure 1, Supplementary Figure S1). Our phylogenetic analysis also identified a third and novel gregarine clade (lineage C) that is weakly associated (bootstrap support = 63%) with the group comprising lineages A, B and *N. temporariae*. Sequences from lineage C were only identified in *M. margaritifera* individuals collected from the healthy reference population at Stommebäcken (Figure 1, Supplementary Figure S1).

**Figure 1.**
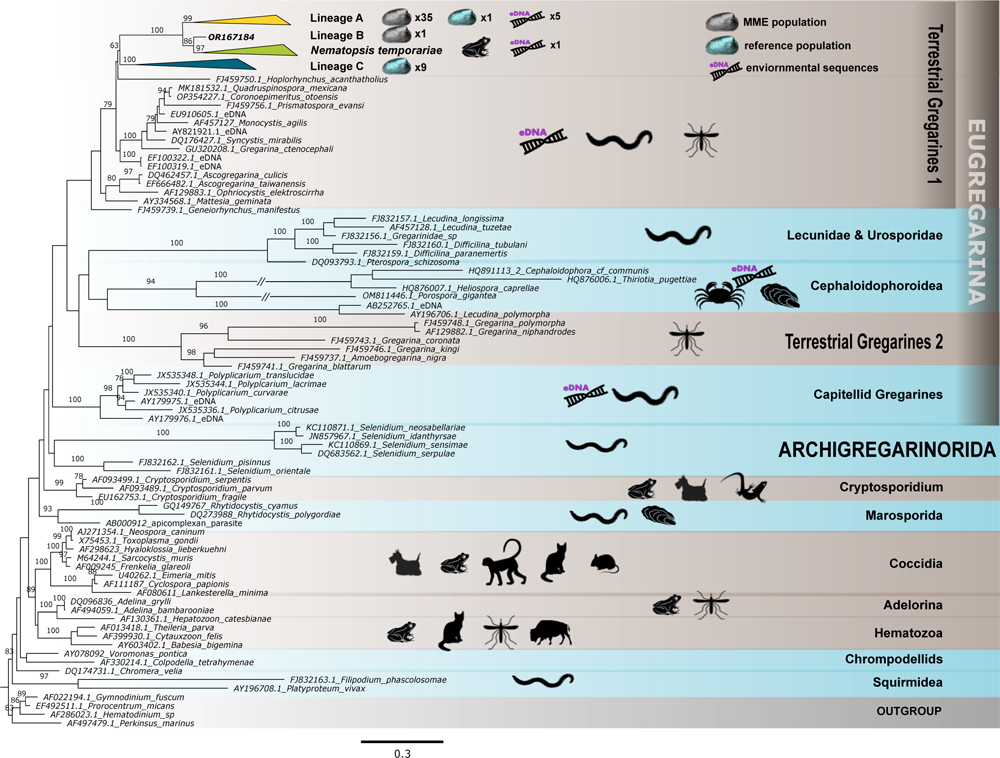
Maximum Likelihood phylogenetic tree based on the 18S rDNA gene with placement of the newly produced gregarine sequences isolated from MME and reference populations of *M. margaritifera.* The analysis was performed with the GTR+F+I+G4 model of evolution. The numbers on branches are bootstrap support after 100 replicates except; bootstrap support below 70% are not shown. For all NCBI sequences, the GenBank accession number and the taxon name are listed. The double slashed line represents manually shortened branches for fitting purposes. Known hosts and environmental sequences are shown using pictograms. The tree was rooted with four sequences belonging to Perkinsozoa and Dinophyceae as outgroups. Lineages A, C and the *N. temporariaea* clade were collapsed for clarity but their full version is available in Figure S1. The scale bar represents the number substitiutions/site.

### Lesions in *M. margaritifera* tissues collected in MME

External inspection of specimens collected in the MME population from Teåkersälven revealed thickening and whitening of the mantel and reductions in gonads (Supplementary Figure S2A). Upon gross morphological investigation, no gonads were visible in the 2018 samples. In 2020, only 1/20 specimens (5%) showed gonads detectable by eye (Supplementary Figure S2B-C). Macroscopic observations of the digestive system, foot, connective tissue and muscle layers did not show any clear signs of discoloration or emaciation (Supplementary Table S2).

Closer histological examinations revealed several lesions in the mantel, gills, digestive glands and gonads. In addition, patterns of increased hemolymph infiltration within connective tissue were found in the mantel (Figure 2A), likely expanding its thickness. The hemolymph regions of the mantel were highly infiltrated by hemocytes in 4/5 mussels collected in September 2018, migrating from blood sinuses into connective tissue and the hemolymph-rich areas (Figure 2B). The hemocytes were hypertrophied or irregularly shaped often with a dislocated nucleus, enclosing weakly stained or brown structures (Supplementary S3A-B). A similar pattern was seen in many hemocytes or phagocytic cells within the auricles of the heart (Figure 2C, Supplementary S3C). These enlarged phagocytic cells reached 11.3 μm (n=4, stdev=0,79) in the mantel and 13.4 μm (n=9, stdev=0.97) in the heart. Cellular infiltration or hyperplasia within gills and mantel was observed forming basophilic clustering of cells within or attached to skeletal parts of the gills as well as in hemolymph gill sinuses and within connective tissue, Figure 2D-F, Supplementary S3D).

**Figure 2.**
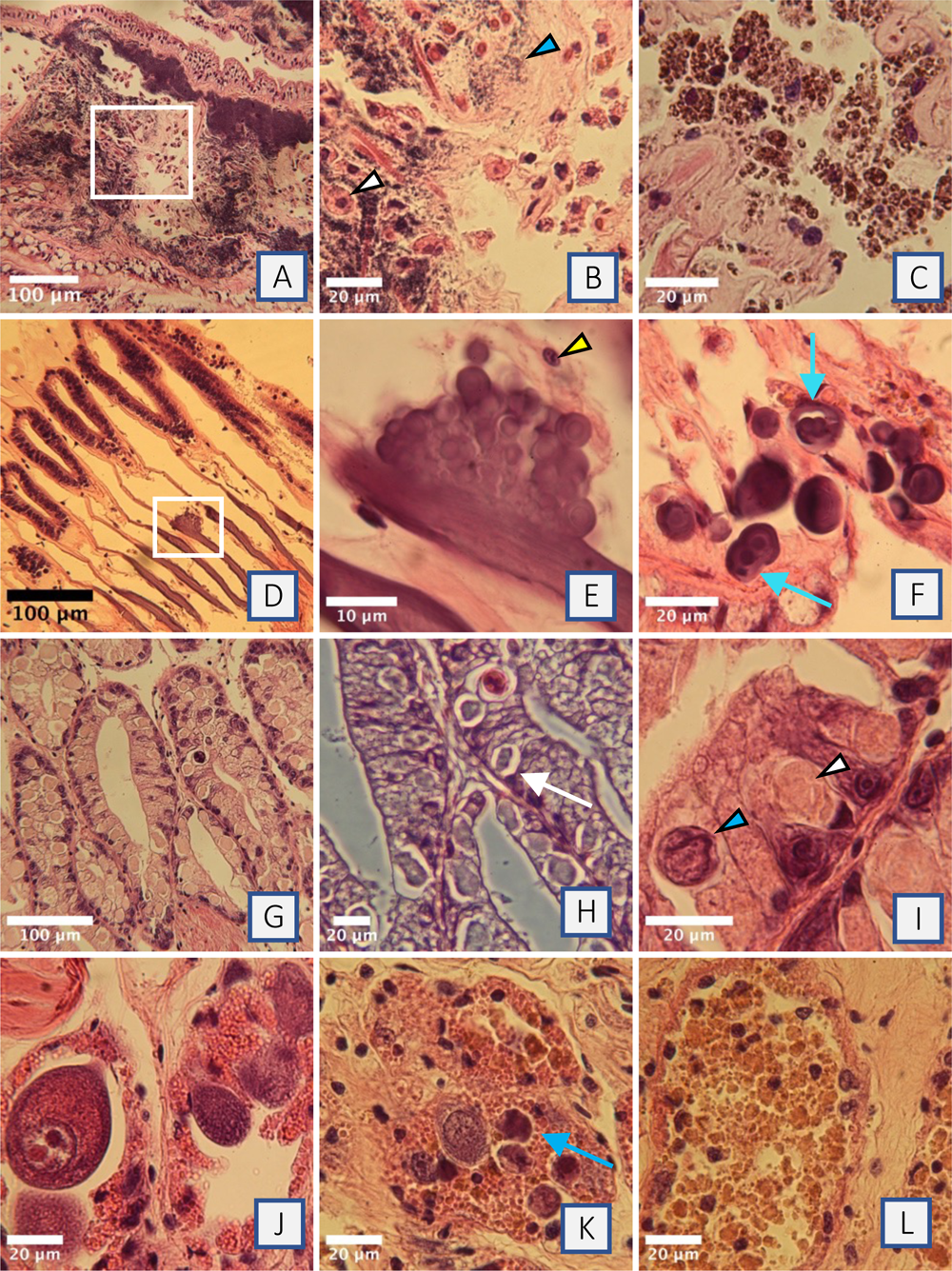
Histopathological lesions associated with gregarine infiltration of mussels collected from the MME population. (A) Infiltration of hemolymph (basophilic droplets) and hemocytes, within the connective tissue of the mantel. (B) Enlarged area (white square), showing diffuse hemolymph spread within connective tissue (blue arrowhead) and invasion of hemocytes (white arrowhead). Note that some hemocyte-nuclei are displaced indicating phagocytized material within cytoplasm. (C) Large phagocytic cells within heart auricles, enclosing multiple pigmented granules or micro-cells. (D) Clusters of basophilic cells expanding from the gill filaments, chitinous skeleton part. (E) Close up of the hyperplasia (white square), where the basophilic cells are characterized by thick cell-walls developing within host epithelia of the gill. Host nuclei indicated with yellow arrow. (F) Gill vessels or sinuses, infiltrated with large and thick-walled basophilic spheres. Signs of cell division are indicated by blue arrows. (G) Large numbers of vacuoles developing within the digestive gland epithelia, enclosing weakly eosinophilic or basophilic cells. (H) Parasitic vacuoles within the vesicular cells (VC) visualized by phase contrast (arrow). (I) The spherical cells within the vacuoles indicated different origin, enclosing both detachment of basophilic cell (blue arrowhead) and cyst formation within eosinophilic vesicular cells (white arrowhead). (J) Gonad follicles enclosing normal oocytes. (K) Ovary follicles filled with developing oocytes as well as pycnotic and degenerative cells (blue arrowhead). Increased numbers of pigmented granules. (L) Gonad follicle without signs of gonad development. Follicles filled with brown degenerative matter.

Degenerative changes were also observed within the digestive gland, primarily in September 2018. All five mussels collected from the MME population exhibited extensive signs of vacuolization, affecting over 50% of the cells (Figure 2G-H). Epithelial lysis, necrosis or disintegration caused detachment of many cells from basal layers, releasing cells or cell content into the lumen of the digestive glands (Supplementary S3D-H). This affected both vesicular and basophilic cells within tertiary gland ducts (Figure 2I). Minor changes and much less vacuolization were observed in June, where 11 out of 20 mussels showed signs of vacuolization and in much lower numbers (less than 10 cells/section). Finally, most gonads exhibited impaired development with reduced or absence of normal cells. In the MME group, only one specimen had regular oocytes (Figure 2J), while many ovaries showed cells with signs of atrophy, where many appeared pyknotic and became densely basophilic (Figure 1K). Among mussels collected in June, 17 out of 20 specimens had completely depleted gonad cells, with follicles containing cellular debris instead (Figure 2L).

### Putative infiltration of a new gregarine parasite

To explore the potential link between the lesions described above caused by invading cells in different tissues and the new sequences obtained, *in situ* hybridization (ISH) was utilized with probes specifically designed to target the isolates of lineage A. Mussels collected in 2018 and 2020 in Teåkersälven (two specimens in each years) revealed gregarine cells in the hemolymph compartment and gills as well as within digestive gland. Mono or dizoic cysts were observed in hemolymph sinuses in the mantel (Figure 3A-B). In the skeletal part of the gill, zoites were forming mono-, di-or polyzoic cysts, and were attached to the endothelium of the gill vessels (Figure 3C-E). A systemic spread and low numbers of monozoic thick walled, histozoic sporocysts were detected embedded in other organs such as digestive gland (Figure 3F). Free zoites were also detected within the lumen of the tertiary digestive glands (Figure 3G), indicating oral uptake of detached cells from the mantel and gills (Supplementary Figure S3E). Within digestive glands, invading zoites were observed attaching to the surface of epithelial cells of tubuli, as well as proliferating within the tubular epithelia, migrating deep within the digestive gland. These infiltrating zoites gave a strong positive ISH staining in June (Figure 3H-I), but larger sporoblast containing immature oocyst stages within the digestive gland gave a weaker signal (Figure 2J-K), and only weak or no signal were recovered from September.

**Figure 3.**
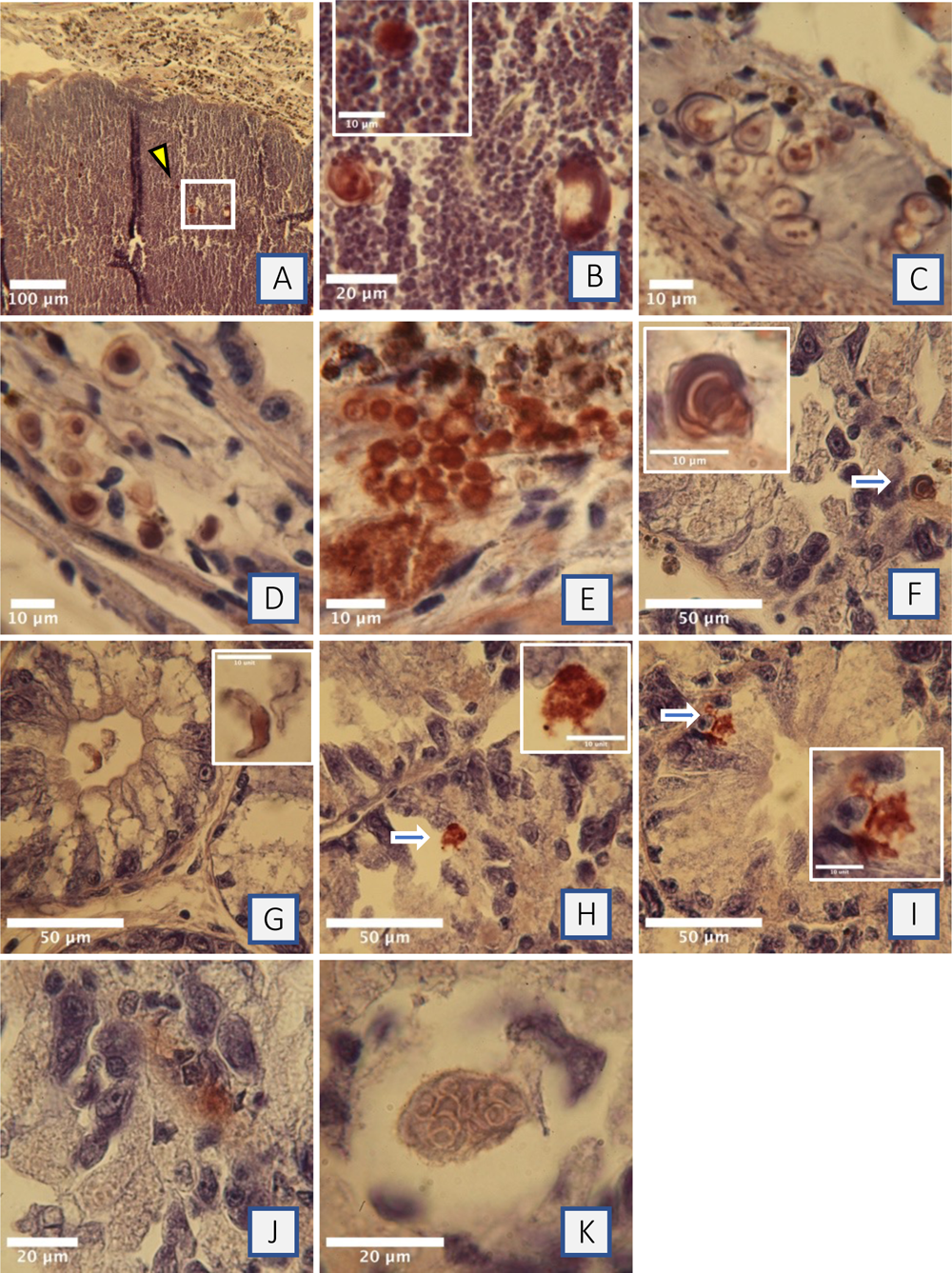
In situ hybridization of lineage A cells showing different development stages in *M. margaritifera* from Teåkersälven. Images A-D correspond to specimens collected in September 2018, images E-K correspond to specimens collected in June 2020. (A-B) Positive reaction (Nova Red, avidin biotin complex) in dorsal hemolymph sinuses, adjacent to heart auricles, visualizing circulating zoites. (C) Developing zoites embedded within gill skeleton parts. (D-E) Gill sections showing multiple crescent folded zoites infiltrating epithelial layers. (F) Section of digestive glands showing a single histozoic cyst embedded between the digestive glands, adjacent or within capillary vessels. (G) Infiltrating zoites within lumen of digestive glands. (H-I) Invasion of zoites attaching to and infiltrating vesicular cells. (J) Early sporogony development, zygote or sporoblast deeply embedded within digestive gland cells. (K) Multicellular sporoblast, within parasitic vacuole, enclosing multiple sporogonic developmental stages.

### Developmental forms of the new parasite

Light microscopy revealed eosinophilic but weakly stained zoites within the hemocytes circulating through the mantel, gills, and heart, sometimes dislocating the host cell nucleus (Figure 4A-B, Supplementary S4 A-B). Large numbers of small round or vermiform cells were also seen within hemocytes in the heart’s auricles, presumably due to phagocytosis (Figure 4C, Supplementary Figure S3C). These cells might correspond to invading naked gregarine infiltrating cells, e.g. similar to gymnospores known to infect marine bivalves (Prytherch 1940). The intracellular circular spores exhibited considerable variation in size and shape, with a mean diameter of 3.06 μm (n=20, stdev=0.75).

**Figure 4.**
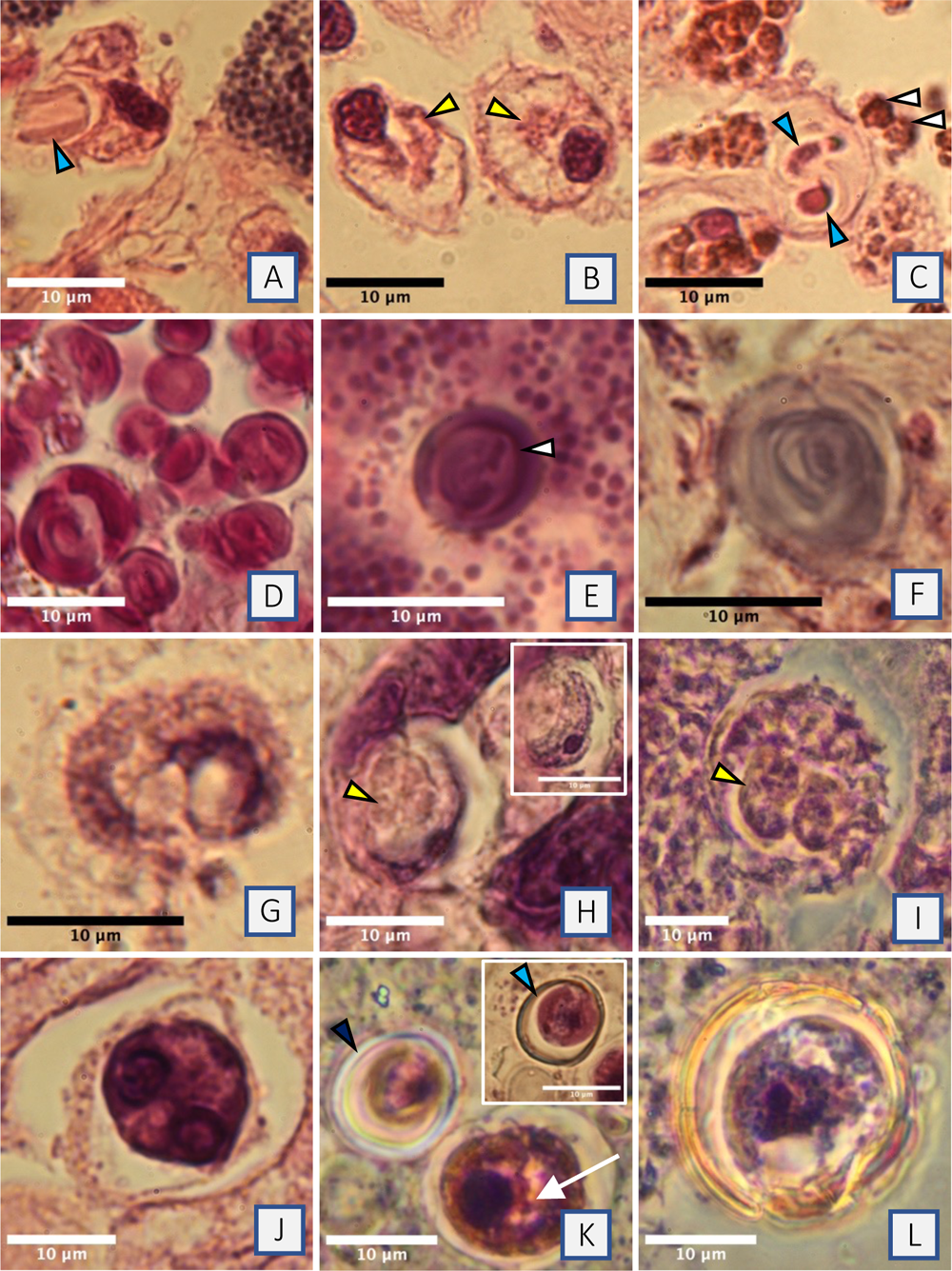
Histological sections of *M. margaritifera*, with morphological descriptions of gregarine cells within different organs. (A) Hemocyte phagocytizing or releasing a folded zoite enclosed within a vacuole. (B) Section of heart auricles indicating infiltration by zoites within the cytoplasm of hemocytes, indicated with yellow arrowheads. (C) Early formation of gregarine cyst after infiltration of connective tissue within heart auricles (blue arrowhead). Adjacent, gymnospore-like cells, indicated with white arrowhead. (D) Hemolymph compartment with development of multiple zoites enclosed within droplets (i.e., naked gregarine spores, gymnospores). (E) Thick-walled gregarine cyst within hemolymph compartment. A single basophilic zoite clearly seen within the cyst wall. (F) Histozoic gregarine spore formation, weakly basophilic, within connective tissue of heart auricles. (G) Detaching spore / zoite, released from the epithelia layers of mantel. (H) section of digestive gland with sporoblast formation attached to basophilic cell nuclei enclosed within a parasitic vacuole. Enclosed formations of internal cysts seen in the center of the sproblast (yellow arrowhead, phase contrast). (I) Early oocyst development. partly enclosing internal sporocysts or immature of sporozoite, phase contrast. (J) Digestive gland with one basophilic sporulated cyst enclosed within residual external membrane and still enclosed in a parasitic vacuole. Within the cyst, two folded sporozoites (darkly basophilic vermiform cells) can be seen. (K) Section of intestine revealing developing sporocysts with double layered soft cyst-wall (blue arrowhead. The same cell indicted, black arrowhead by phase contrast). The larger cyst is unsporulated with signs of further separation into single sporocyst indicated by white arrow. (L) Large sporocyst enclosing single sporozoite. The external cystwall were refractile/weakly yellow visualized by phase contrast, indicating thick and rigid cystwall formation. This may indicate a fully sporulated cyst before released as fecal outlet.

The hemolymph was distributed as strongly basophilic droplets, which were infiltrated with polymorphic zoites enclosed in transparent spheres in both the mantel and gills. Single or double vermiform stages, measuring approximately 2-15 μm in length, were folded into open semicircles or closing full circles within these spheres (Figure 4D, Supplementary S4D-E). In addition, mono/dizoic thick walled cysts were enclosed in connective tissue, as histozoic cysts of the gills and mantel as well as freely dispursed within the hemolymph compartment (Figure 2E-F, 4E-F, Supplementary S4F), morphologically corresponding to the cells observed by ISH (Figure 3A-E). Many cysts enclosed more than one zoite, indicating proliferating cells, forming two or multiple zoites within the same oocyst (Supplementary S3D, S4G). The examination of the mantel and gills revealed signs of epithelial loss, as well as release of hemocytes and sporoblast-like cells (Figure 4G). Similar cells were observed infiltrating the digestive gland indicating an oral route via the stomach and further to the tubuli channels. Multiple such cells were seen entering the digestive glands through digestive tubules (Supplementary Figure S4H), an observation also supported by ISH (Figure 3G).

The pattern of vacuoles forming within the digestive gland of the mussels from MME population was further investigated by light microscopy. Phase contrast showed formation of parasitophorous vacuoles where pairs of zoites (presumably zygote or syzygy formation) developed into sporoblasts enclosing oocysts with inner sporocysts (Figure 4H, I). These oocysts, reaching over 20 μm in size (Figure 4I), enclosed two inner sporocysts whithout clear signs of inner sporozoites, i.e., unsporulates oocysts. Two types of larger vermiform parasite cells, basophilic and eosinophilic, were also observed in front to front position, which we interpret as merging gamonts in a process termed syzygy (Supplementary Figure S4I), indicating a gregarine gamogonial phase. From ruptured parts of the digestive glands, immature sporocysts as well as mature disporous ones were also found (Figure 4J, Supplementary Figure S4J-K).

The majority of the oocysts were released unsporulated, thus the intestine was also investigated to follow potential release of parasites as fecal outlet. The intestinal lumen contained multiple sporocysts, suggesting a link between immature sporozoites (sporoblasts) and oocysts that could have developed into free sporocysts during the release. These undifferentiated sporocysts with a mean diameter of 14.5×13.9 μm (n=5) likely also divided further in two parts as seen on Figure 3K. Two types of cysts showed large variation in size and shape: smaller cysts (Figure 4K) with a mean diameter of 9.75×9.0 μm (n=22, range 6.8-11.2μm), and larger thick walled cysts with a diameter of 21.7 x 21.2 μm (n=1). Since no fecal samples were investigated directly upon sampling, the final morphology of sporulated sporocysts are unresolved, even though the double, thick and refringent walled sporocyst in Figure 3L could represent such a final stage.

### Ultrastructure

Transmission electron microscopy (TEM) showed small zoites with alveoli-and rhoptry-like organelles in the lumen of digestive tubuli (Figure 5A,B). Infiltrating zoites seemed to induce local recruitment of microvilli enclosing them within the host epithelium (Figure 5C), or induce openings between epithelial cells, reaching the basal regions deeply within the digestive tubular cells (Figure 5D). Furthermore, we found indication for sexual replication (gamogony) as two types of gamogonial stages were found within the digestive glands. A free small gamont-like cell without flagellae (Figure 5E,F) and a large gamont-like cells containing an organelle with mitochondria-like morphology, developing within parasitophorous vacuoles (Figure 5G,H). The ultrastructure demonstrated fusion of gamont-cells forming zygotes supporting anisogamogonial cell development as part of the lifecycle (Figure 5I). In semithin sections, signs of gamonts in similar formations adjacent to sporogonic formations were observed within parasitophorous vacuoles (Figure 5J,K).

**Figure 5:**
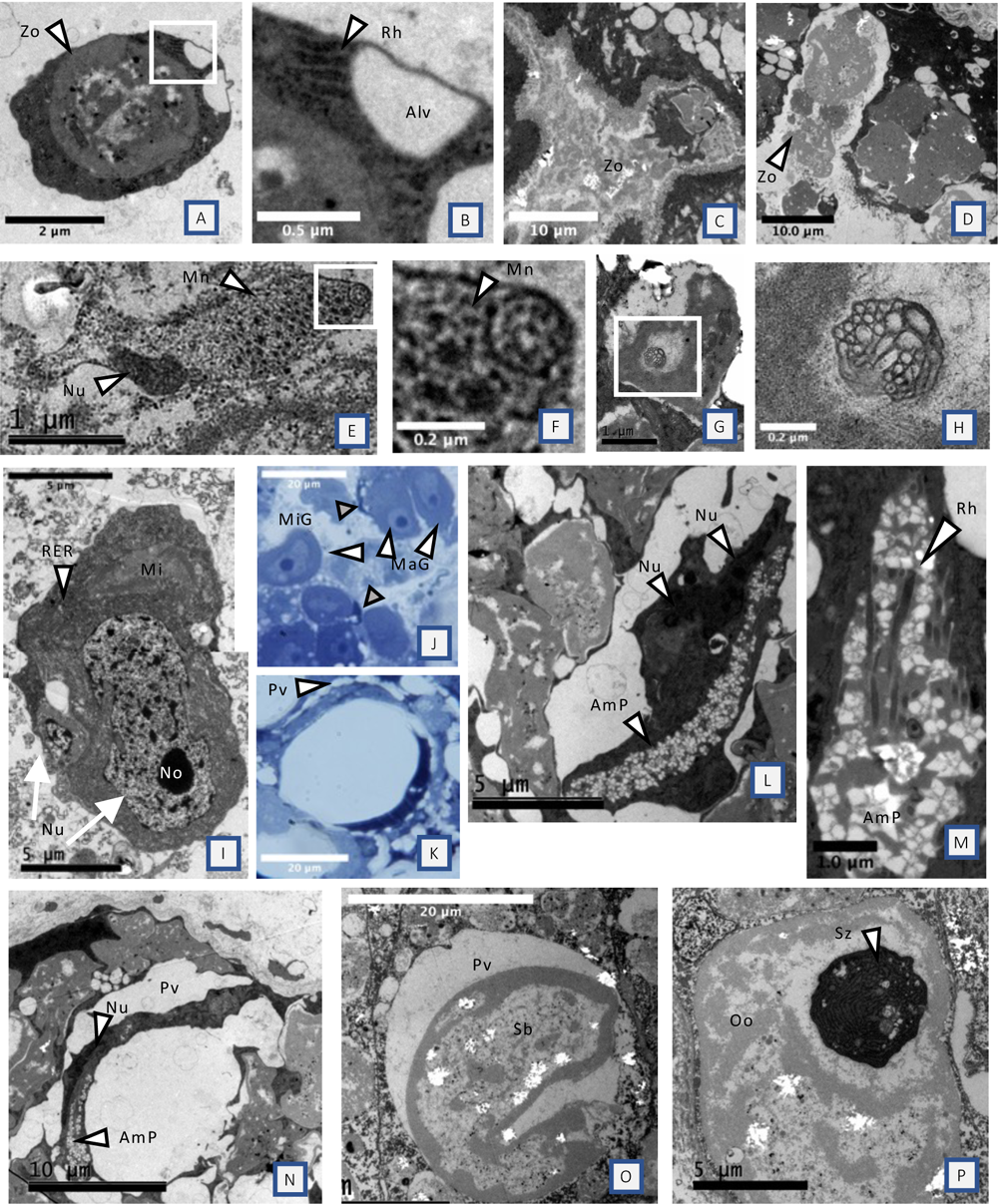
Ultrastructure of digestive gland with micrographs of putative gregarine infiltration and indication of gamogony and sporogony. (A-B) Free zoite within lumen of tertiary digestive gland. Rhophtry and alveoli like organelles indicated. (C) Infiltration of zoite enclosed by host microvilli. (D) Closely associated zoites with larger (electron lucent) and smaller (electron dense) cells arranged in a chain, reaching deep between digestive gland epithelium. (E-F) Free small gamont-like cell, with mitochondria and microneme-like structures. (G-H) Large gamont-like cell within parasitophorous vacuole with clear mitochondria like organelle in the center (enlarged in image H). (I) Fusion of a small and large gamont forming a zygote. Rough endoplasmatic reticulum dominating the cytoplasm. (J) multiple larger (white arrowhead) and smaller (grey arrowhead) gamonts. (K) Formation of a large parasitophorous vacuole, enclosing one darkblue sporozoite and one light blue sporoblast and multiple vacuoles in the external rim (white arrowhead). (L) Zygote (electrondense) with formation of amylopectin granules, arranged in a strings with snowflake like pattern. Fusion of nuclei indicated with arrowhead. (M) Close up visualizing the amylopectin like granules infiltrated with rhoptry like organelles. (N) Formation of a parasitophorous vacuole enclosing a sporozoite forming into a closing circle. Note the nuclei and amylopectin like granule indicated with white arrowheads. (O) Sporoblast (elecronlucent) enclosed within parasitophorous vacuole. (P) Oocyst (electron lucent) with formation of a sporozoite (electron dense) in cross section. J-K: Semithin sections. tuloidin stained. Abbreviations: Zo=zoite, Rh=rhoptry like, Alv=alveoli-like, Nu=nucleus, Mn=microneme, RER=ribosomal endoplasmatic reticulum, MIG=Microgamont, MaG=Macrogamont, AmP=amylopectin like granules, Pv= parasitophorous vacuole, Sb=sporoblast, Oo=oocyst, Sz=sporozoite. Size indicated by scalebar in image. Abbreviation: Zo-Zoite, Rh-rhoptry, Alv-alveoli, Nu-nuclei, No-nucleolus, MaG-large gamont,, MiG-small gamont, AmP-amylopectin like structures, Sb-sporoblast, Oo-Oocyst, Sz_Sporozoit.

Gamonts in zygote formation and sporozoites can form clusters of amylopectin granules (Valigurová and Koudela 2006). Within electrondense vermozoite cells, we detected long clusters of amyloid-pectin-like granules arranged in snowflake pattern, also infiltrated with rhoptry-like structures. The ultrastructure shows zygotes forming into oval semicircles and potentially spherical oocysts (Figure 5L-N). This could be the start of a sporogonial development with the formation of oocysts and sporocysts within digestive glands. Multiple electron lucent sporoblasts or immature oocysts (Figure 5O) were proliferating in large numbers primarily in the mussels collected in September, often enclosed within a vacuole. Within these, a few oocysts also developed inner electron dense sporocysts or enclosed sporozoites that appeared almost black under electron microscopy (Figure 5P).

## DISCUSSION

We report the findings of novel diversity of gregarines associated with the highly endangered fresh water perl mussel species *M. margaritifera* in Swedish streams. Two main new groups of related 18S rDNA sequences were obtained (Figure 1). One group (lineage A) is related to a described parasite in tadpoles, *Nematopsis temporariae*, and was found almost exclusively (with one exception – sequence accession number: OR168627) in association with individuals suffering mass mortality events (Figure 1, Supplementary Figure S1 and Table S4. Five environmental sequences from lakes and permafrost soil also belong to this group. The other main group (lineage C) was only recovered from *M. margaritifera* collected in the reference population (Stommebäcken), which were not experiencing mass mortality at the time of sampling and did not show signs of a gregarine infection. Further efforts are needed to understand the nature of host interaction of these gregarine species (Rueckert, Betts, and Tsaousis 2019). In addition to these two main groups, we also recovered in the MME population one sequence (lineage B), which is most closely related to the *N. temporariae* parasites found in tadpoles (Chambouvet et al. 2016).

The genus *Nematopsis* has long been known since the first morphological description of a parasitic infection in scallops in 1892 (Kinne and Lauckner 1983). Since then, many reports have been published of cells attributed to *Nematopsis,* mainly in marine invertebrates such as crustaceans and bivalves, all based on morphological descriptions of gamogony and sporogony (bivalve hosts), and schizogony (crustacean hosts) (Chakraborti and Bandyopadhyay 2010; Silva et al. 2019; Abdel-Baki et al. 2012; C. Azevedo and Cachola 1992; Vivares 1969). The *Nematopsis* species known to infect marine bivalves are characterized by monozoic, naked sporozoites within thick walled cysts (Kinne and Lauckner 1983). *Nematopsis spp.* is an exception among gregarines as they typically use two hosts to complete their lifecycle. Recent investigations indicated that the known host range of *Nematopsis* species also includes marine polychaetes (Morales-Covarrubias et al. 2014) and, importantly, tadpoles in freshwater environment (Chambouvet et al. 2016).

The tadpole parasite (*N. temporariae*) is the first *Nematopsis* species with both morphology and molecular data available. However, the assignment of this parasite to *Nematopsis* and its phylogenetic placement in a terrestrial clade challenged previous taxonomy based on morphology, which has considered *Nematopsis* as part of the marine clade Porosporidae (Cavalier-Smith 2014). This is further conflicting with a recent study placing *Porosporida gigantea* as sister group to marine gregarine and not with *N. temporariae* (Boisard et 2022). Thus, it is currently uncertain whether *N. temporariae* and the new associated diversity we describe here are truly members of the genus *Nematopsis*. Clarification of the affiliation of these sequences will require obtaining molecular data for marine *Nematopsis*.

The possibility of a wrong assignment to *Nematopsis* of the tadpole parasite is supported by our microscopic investigations of the new parasitic diversity within *M. margaritifera*, which revealed some similarities but also key differences with published reports of marine *Nematopsis*. For example, clusters of cysts in gills containing single or multiple vermozoite-like cells (Figure 1) are similar to descriptions of *Nematopsis* cysts in marine bivalves such as scallops, blue mussels and oysters (Kinne & Lauckner, 1983; Prytherch, 1940; Vivares, 1969). Within the digestive system, a few darkly eosinophilic cysts were detected (Figure 3J), again similar to cysts described in marine *Nematopsis* (Belafastova, 1996; Kinne & Lauckner, 1983; Prytherch, 1940; Silva et al., 2019; Tuntiwaranuruk et al., 2004; Vivares, 1969). This was also the case of the single sporozoite observed within monozoic sporocysts (Figure 3L), corresponding to be the final stage of *Nemotopsis* cyst formation in marine molluscs (Prytherch, 1940). However, all marine *Nematopsis* descriptions refer to histozoic cyst formation, enclosing cysts within connective tissue (Kinne and Lauckner 1983; Vivares 1969; Silva et al. 2019; Tuntiwaranuruk et al. 2004). Our observations partly support the presence of such histozoic cysts, but the ongoing oocyst release from digestive gland and transformation into sporocysts later via intestine passage as fecal release (Figure 3K-L) do not correspond to these descriptions. Furthermore, the transformation of oocysts into sporocysts enclosing double sporozoites and further separation into monozoic sporocysts differ from the descriptions of marine *Nematopsis spp.*, where clusters or single gregarine always sporulates inside an organ without release. Description of the tadpole infection in liver also indicates that these gregarine cysts are released from the host, since only empty oocysts could be recovered after the metamorphose phase, when tadpoles have transformed into subadult frogs (Chambouvet et al. 2016). These reported gregarine cysts within tadpole livers appear not thickwalled and thus do not fully correspond to the morphological descriptions of marine *Nematopsis* were the thick cystwall is a defining feature of this genus (Kinne and Lauckner 1983).

Based on our observations across various organs we suggest a putative life cycle for the new gregarine parasite (lineage A) found in *M. margarifiera* (Figure 6). Very few life cycle descriptions exist for gregarines in bivalves, all based on marine hosts, so we have used the description of *Nematopsis ostrearum* for comparison (Prytherch 1940). We hypothetise that cell infiltration begins within the epithelium of gills and mantle by infiltration of naked spores (gymnospores) where development starts (asexual replication), which then spreads throughout the host via hemolymph or circulating hemocytes (Figure 6A-D). Further assexual division results in mono/di/polyzoic cysts that invade the connective tissue of different organs as histozoic cysts. Since only very few of these were detected embedded within connective tissue of the digestive glands, the proliferation within digestive tubules is more likely distributed within the host as oral uptake. Zoites released from initial cell division in the gills and mantle are therefore the more likely transport route, leading cells into the stomach and further to the digestive gland via the digestive ducts (Figure 6F-G). These parasitic cells infiltrate the epithelia of the digestive cells, where cell division continues (Figure 6H). Paring of zoites involving larger and smaller elongated cells in syzygy-like position was observed within digestive tubuli (Figure 6I-J) supporting gamogonial development. The fusion of gametes into zygotes and further into sporulating cysts (oocysts), would represent a sporogonic part of the lifecycle (Figure 6K).

**Figure 6.**
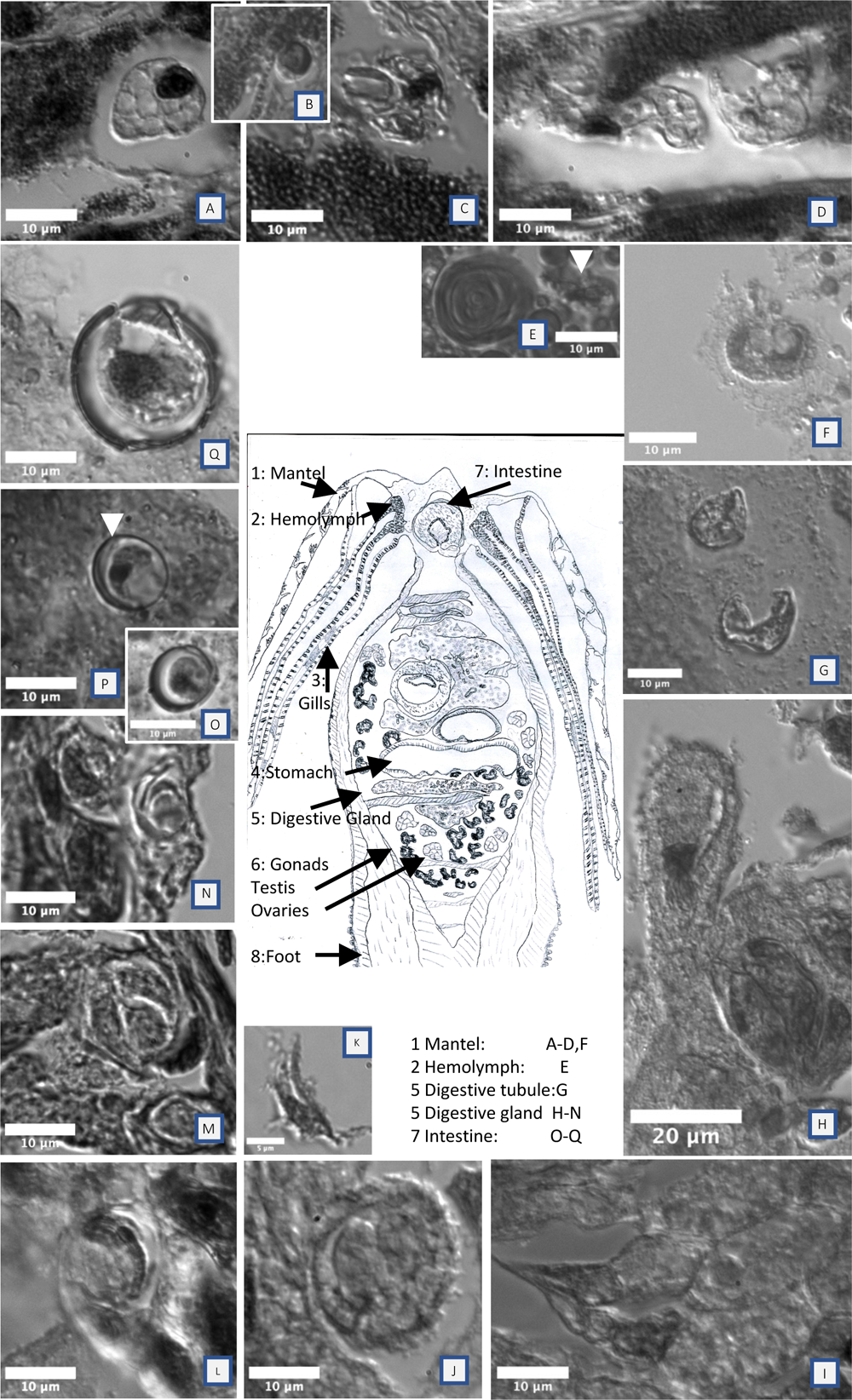
Hypothetical lifecycle and observations of the new gregarine lineage A within the host *M. margaritifera* from Teåkersälven (MME population). (A) Sporozoites or gymnospores from the environment are likely infiltrating gills and mantel via host feeding activity e.g., water filtration of food particles and further taken up via host cell infiltration or phagocytic activity of hemocytes and epithelial cells. (B) Phagocytized cells expand in size and (C) are released from rupturing host cells to develop and (D) infiltrate other host cells. (E) Further asexual division results in mono/di/polyzoic cysts that invade the connective tissue as histozoic cysts across different organs, transported via hemolymph or circulating hemocytes. (F) Gregarine zoites or histozoic cysts within the mantel and gills were also observed, released from rupturing epithelia of host cell. (G) These released zoites are likely transported as food particles to the stomach and into the digestive glands via the digestive ducts and (H) where zoites attach and become epicellularly enclosed within at the epithelia of digestive tubules. Many zoites were observed inside host cells enclosed within a parasitophorous vacuole. (I) Paring of zoites involving larger elongated cells in syzygy-like position, was detected within digestive tubule and (J) enclosed into cysts supporting gamogonial development. (K, L) The fusion of gametes into zygotes and further into sporulating cysts (oocysts) may represent a sporogonic part of the lifecycle. (O-Q) Very few mature oocysts were detected within the digestive glands indicating sporulation during fecal release, also supported from observations of the intestinal lumen where multiple monozoic oocysts also was observed. Schematic figure of crosssection of *M. margaririfera* in the center, indicates anatomical organization of the host, to further illustrate the gregarine infiltration.

Within the digestive gland, sporogony are likely the start of new proliferative cycles were fusing gametes produce sporoblasts. These developed further into immature oocysts that grow and expand in size, enclosing two or four sporocysts/sporozoites within the oocyst (Figure 6L), similar to descriptions of eugregarine sporogony in insects (Dias et al. 2017). In September, large numbers of parasitic vacuoles were observed contaning single to multiple unsporulated oocysts, which were shed via rupturing of the digestive gland epithelium (Figure 6M-N). In the intestine, further maturation of unsporulated oocysts were observed, producing mono-or dizoic sporocysts. However, the formation of different sized sporocyst cells was puzzling, with smaller and possibly double overlapping elastic cystwalls walls (Figure 6O-P), and few examples of larger thick walled refringent and dense cystwall (Figure 6Q) indicating potentially two types of sporocysts. The final morphology with fully sporulated cysts is therefore still unclear.

All known eugregarine infections are monocyclic and often develops in the lumen of digestive system within insect, normally attached to epithelial cells (Clopton 2002). However, there are exceptions such as some ascogregarine infections of mosquitos where the parasites develop intracellularly within epithelia cells and are shed as oocysts via fecal release (Lantova and Volf 2014), similar to the putative life cycle presented here. Other gregarine-related parasites infecting aquatic hosts, such as *Cryptosporidium* spp. found in fish, also exemplify intracellular cell development in which the sporogonial part takes place in parasitophorous vacuoles deeply nested within the cytoplasm of the epithelial cells, in contrast to all other cryptosporidians in vertebrates where development takes place in apical position attached to the epithelial cells (Alvarez-Pellitero and Sitjà-Bobadilla 2002; Couso-Pérez, Ares-Mazás, and Gómez-Couso 2022).

A lot remain to be understood about the novel gregarine diversity we report here and its interaction with *M. margaritifera*. Most importantly, the significance of the observed summer seasonality of lineage A in the mussel host is unclear, but could indicate that other hosts are involved in other parts of the years. This would be a novelty as all descriptions of terrestrial gregarines thus far support monoxeneus life cycles. An in-depth description of the life stages of the novel lineage C within *M. margaritifera*, and its impact on the host, also requires further investigations. But our work clearly reveals that a diversity of gregarine parasites is found within Freshwater perl mussels *M. margaritifera*, some associated with mass mortality of this ecologically critical species in streams and rivers. In future, this knowledge should be taken into account in conservation effort of this endangered species.

## Acknowedgements

This research was supported by SciLifeLab, a Formas Future Research Leader grant (2017-01197) and a Swedish Research Council VR research project grant (2017-04563) both available to FB

We are thankful to Västra Götaland county, Dalslands miljö och energiförbund, and the Swedish agency for Marine and Water Management for funding the collection and preparation of freshwater pearl mussels. We thank Ulrika Larsson Pettersson at the Department of Pathology and Wildlife (Swedish National Veterinary Institute) for assistance with ISH, and Karin Staxäng at the Biovis platform (Uppsala University) for TEM-preparation and guidance in TEM imaging. Finally, we are grateful to Mark Freeman at Ross University School of Veterinary Medicine (West Indies) for help with ISH probe design. Computational analyses were enabled by resources in projects SNIC2021/5-302, SNIC2021/22-178, NAISS2023-6-81and NAISS2023-5-122 provided by the Swedish National Infrastructure for Computing (SNIC) at UPPMAX, partially funded by the Swedish Research Council through grant agreement number 2018-0597.

## Author Contributions

AA and FB designed the study. AA and NW performed field and sample collection. AA performed lab work. AA, IOB, NW, MJ, AK analysed data and prepared Figures. AA, IOB, FB, DP, AK and MJ wrote and edited the paper.

