## Supplementary tables and figures for "Identification of a new gregarine parasite in mass mortality events of freshwater pearl mussels (*Margaritifera margaritifera*) in Sweden"

**Supplementary Table S1**. Field sampling of freshwater pearl mussels (*Margaritifera margaritifera*) with population data from Stommebäcken (reference population) and Teåkersälven with MME's. The population in Stommebäcken is used as a control as it has not been detected any mass mortality events during the investigations. The population in Teåkersälven have been investigated seven times between 1985 and 2019 and there was a dramatic decline between 2016-2019 caused by a mass mortality event that was detected in 2017 (Wengström et al., 2019).

| **Stream** | **Survey year** | **Population size** | **% juveniles (<50mm)** |
|---|---|---|---|
| Stommebäcken | 1990 | 2130 | 11 |
| Stommebäcken | 1997 | 2193 | 2 |
| Stommebäcken | 2004 | 1727 | 0 |
| Stommebäcken | 2010 | 2610 | 1 |
| Stommebäcken | 2016 | 2728 | 0 |
| Teåkersälven | 1985 | 84822 | - |
| Teåkersälven | 1990 | 43950 | 0 |
| Teåkersälven | 1997 | 65291 | 1 |
| Teåkersälven | 2003 | 46939 | 0 |
| Teåkersälven | 2010 | 49336 | 0 |
| Teåkersälven | 2016 | 45098 | 0 |
| Teåkersälven | 2019 | 15774 | 0 |

Live specimens of freshwater pearl mussels were collected from both streams by wading with a bathyscope searching the bottom for mussels. In Teåkersälven the aim was to collect mussels that showed signs of sickness but that were still alive, e.g., laying on top of the substrate, gaping and slow to respond to tactile stimuli (Richard *et al.*, 2020). In Stommebäcken we collected mussels from an area with a dense mussel bed (>1 mussel/$m^2$) and on this site there where no signs of sick mussels.

**Supplementary Table S2**. Biological parameters and macroscopical findings from examination of freshwater pearl mussels (*Margaritifera margaritifera*). Stommebäcken unaffected streams (reference population) and Teåkersälven with MME's. Most prominent findings were that gonads were not detected in many individual and not only affecting the MME population but also detected in the reference population indicating an impact on reproduction capability affecting both populations. Missing data marked with -. Macroscopic examination not applicable for gonads, foot and digestive gland due to fixation in field. Investigations follow in line with previous work (Wengström et. al. 2019).

| STREAM NAME | SURVEY YEAR | DATE | ID | LENGTH (CM) | WEIGHT WHOLE SPECMEN | WEIGHT (SOFT BODY) | GONADS (SIZE) | FOOT | DIGESTIVE GLAND | SHELL EXTERNAL/ INTERNAL |
|---|---|---|---|---|---|---|---|---|---|---|
| County of Västra Götaland | | | | | | | | | | |
| Teåkersälven downstream | 2018 | 25 sept | 2599-18 | 10,5 | 63,2 | 19,0 | Not detected | Compact halfway released | Pale green, no content | -/ iridescent |
| Teåkersälven downstream | 2018 | 25 sept | 2606-18 | 10,5 | 73,4 | 21,2 | Not detected | Compact withdrawn | Medium green, no content | -/ iridescent |
| Teåkersälven downstream | 2018 | 25 sept | 2607-18 | 11,5 | 100,8 | 25,4 | Not detected | Compact withdrawn | Light green, no content | -/ iridescent |
| Teåkersälven downstream | 2018 | 25 sept | 2608-18 | 10,2 | 64,4 | 18,2 | Not detected | Compact withdrawn | Dark green, no content | -/ iridescent |
| Teåkersälven downstream | 2018 | 25 sept | 2609-18 | 9,5 | 65,1 | 17,0 | Not detected | Compact withdrawn | Dark green, no content | -/ iridescent |
| Stommebäcken | 2018 | 25 sept | 2610-18 | 8,8 | 37,8 | 10,8 | Not detected | Compact withdrawn | Dark green, no content | -/ iridescent |
| Stommebäcken | 2018 | 25 sept | 2611-18 | 9,0 | 41,5 | 12,0 | Not detected | Compact withdrawn | Light green, no content | -/ iridescent |
| Stommebäcken | 2018 | 25 sept | 2612-18 | 8,2 | 38,8 | 9,9 | Not detected | Compact withdrawn | Medium green, no content | -/ iridescent |
| Teåkersälven downstream | 2019 | 9 oct | 1198-20 | 11,3 | 78,1 | 13,6 | (EtOH) | (EtOH) | (EtOH) | -/ iridescent |
| Teåkersälven downstream | 2019 | 9 oct | 1199-20 | 11,2 | 78,5 | 12,4 | (EtOH) | (EtOH) | (EtOH) | -/ iridescent |
| Teåkersälven downstream | 2019 | 9 oct | 1200-20 | 11,2 | 69,9 | 11,5 | (EtOH) | (EtOH) | (EtOH) | -/ iridescent |
| Teåkersälven downstream | 2019 | 9 oct | 1201-20 | 11,5 | 75,9 | 11,3 | (EtOH) | (EtOH) | (EtOH) | -/ iridescent |
| Teåkersälven downstream | 2019 | 9 oct | 1202-20 | 11,7 | 78,0 | 11,8 | (EtOH) | (EtOH) | (EtOH) | -/ iridescent |
| Teåkersälven downstream | 2019 | 9 oct | 1203-20 | 11,0 | 68,8 | 11,8 | (EtOH) | (EtOH) | (EtOH) | -/ iridescent |
| Teåkersälven downstream | 2019 | 9 oct | 1204-20 | 11,5 | 81,4 | 12,5 | (EtOH) | (EtOH) | (EtOH) | -/ iridescent |
| Teåkersälven downstream | 2019 | 9 oct | 1205-20 | 10,9 | 65,4 | 10,7 | (EtOH) | (EtOH) | (EtOH) | -/ iridescent |
| Teåkersälven downstream | 2019 | 9 oct | 1206-20 | 10,0 | 44,5 | 12,1 | (EtOH) | (EtOH) | (EtOH) | -/ iridescent |
| Teåkersälven downstream | 2019 | 9 oct | 1207-20 | 11,3 | 80,7 | 14,0 | (EtOH) | (EtOH) | (EtOH) | -/ iridescent |
| Teåkersälven downstream | 2020 | 11 june | 2299-20 | 10,7 | 76,6 | 23,3 | Not detected | Compact withdrawn | Medium green, no content | Thick filamented/ iridescent |
| Teåkersälven downstream | 2020 | 11 june | 2300-20 | - | 49,5 | 16,5 | Medium, protruding | Compact withdrawn | Dark green, no content | Thick filamented/ iridescent |
| Teåkersälven downstream | 2020 | 11 june | 2301-20 | 11,4 | 95,4 | 26,2 | Not detected | Compact withdrawn | Dark green, no content | Thick filamented/ iridiscent |
| Teåkersälven downstream | 2020 | 11 june | 2302-20 | 12,3 | 105,6 | 27,5 | Not detected | Relaxed, halfway released | Medium green, no content | Partial filamented/ iridescent |
| Teåkersälven downstream | 2020 | 11 june | 2303-20 | 11,5 | 95,4 | 21,9 | Not detected | Compact withdrawn | Medium green, no content | Thick filamented/ iridescent |
| Teåkersälven downstream | 2020 | 11 june | 2304-20 | 12,0 | 97,8 | 20,4 | Not detected | Compact withdrawn | Pale green, no content | Partial filamented/ iridescent |
| Teåkersälven downstream | 2020 | 11 june | 2305-20 | 11,5 | 91,4 | 26,3 | Not detected | Compact withdrawn | Dark green, no content | Short filamented/ iridescent |
| Teåkersälven downstream | 2020 | 11 june | 2306-20 | 11,7 | 101,9 | 27,7 | Not detected | Compact withdrawn | Dark green, no content | Thick filamented/ iridescent |

| | | | | | | | | | | |
|---|---|---|---|---|---|---|---|---|---|---|
| Teåkersälven downstream | 2020 | 11 june | 2307-20 | 92,4 | 11,8 | 22,6 | Not detected | Relaxed, halfway released | Pale green, no content | Partly filamented/ iridescent |
| Teåkersälven downstream | 2020 | 11 june | 2308-20 | 11,3 | 89,3 | 28,1 | Not detected | Compact withdrawn | Medium green, no content | Thick filamented/ iridescent |
| Teåkersälven upstream | 2020 | 11 june | 2309-20 | 11,0 | 109,7 | 21,1 | Not detected | Relaxed, halfway released | Meidum green, no content | Non filamented/ iridescent |
| Teåkersälven upstream | 2020 | 11 june | 2310-20 | 11,8 | 86,8 | 20,9 | Not detected | Relaxed, fully released | Pale green, no content | Non filamented/ iridescent |
| Teåkersälven upstream | 2020 | 11 june | 2311-20 | 12,0 | 104,7 | 25,1 | Not detected | Relaxed, fully released | Pale green, no content | Non filamented/ iridescent |
| Teåkersälven upstream | 2020 | 11 june | 2312-20 | 11,7 | 84,2 | 20,0 | Not detected | Relaxed, partly released | Pale green, no content | Non filamented/ iridescent |
| Teåkersälven upstream | 2020 | 11 june | 2313-20 | 11,2 | 85,9 | 21,9 | Not detected | Compact withdrawn | Medium green, no content | Non filamented/ iridescent |
| Teåkersälven upstream | 2020 | 11 june | 2314-20 | - | 79,2 | 18,4 | Not detected | Compact withdrawn | Medium green, no content | Non filamented/ iridescent |
| Teåkersälven upstream | 2020 | 11 june | 2315-20 | 11,0 | 86,4 | 21,8 | Not detected | Relaxed, partly released | Pale green, no content | Non filamented/ iridescent |
| Teåkersälven upstream | 2020 | 11 june | 2316-20 | 11,4 | 84,1 | 18,6 | Not detected | Compact withdrawn | Medium green, no content | Non filamented/ iridescent |
| Teåkersälven upstream | 2020 | 11 june | 2317-20 | 11,3 | 90,0 | 20,1 | Not detected | Compact withdrawn | Dark green, no content | Non filamented/ iridescent |
| Teåkersälven upstream | 2020 | 11 june | 2318-20 | 11,3 | 100,6 | 23,3 | Not detected | Compact withdrawn | Dark green, no content | Thick filamented/ iridescent |
| Stommebäcken | 2020 | 11 june | 2319-20 | 10,5 | 58,9 | 14,7 | Not detected | Compact withdrawn | Medium green, no content | Thick filamented/ iridescent |
| Stommebäcken | 2020 | 11 june | 2320-20 | 10,2 | 66,5 | 15,3 | Small | Compact withdrawn | Medium green, no content | Thick filamented/ iridescent |
| Stommebäcken | 2020 | 11 june | 2321-20 | 9,7 | 52,1 | 14,5 | Small | Compact withdrawn | Medium green, no content | Thick filamented/ iridescent |
| Stommebäcken | 2020 | 11 june | 2322-20 | 9,6 | 51,7 | 14,9 | Not detected | Compact withdrawn | Medium green, no content | Thick filamented/ iridescent |
| Stommebäcken | 2020 | 11 june | 2323-20 | 10,6 | 72,7 | 17,9 | Not detected | Compact withdrawn | Dark green, no content | Thick filamented/ iridescent |
| Stommebäcken | 2020 | 11 june | 2324-20 | 9,8 | 49,6 | 15,3 | Small | Compact withdrawn | Dark green, no content | Thick filamented/ iridescent |
| Stommebäcken | 2020 | 11 june | 2325-20 | 10,0 | 56,6 | 13,6 | Small | Compact withdrawn | Dark green, no content | Thick filamentous/ Dark nodules |
| Stommebäcken | 2020 | 11 june | 2326-20 | 11,0 | 80,8 | 17,4 | Not detected | Compact withdrawn | Medium green, no content | Thick filamented/ iridescent |
| Stommebäcken | 2020 | 11 june | 2327-20 | 9,8 | 56,2 | 13,5 | Small | Compact withdrawn | Dark green, no content | Thick filamented/ iridescent |
| Stommebäcken | 2020 | 11 june | 2328-20 | 9,6 | 44,9 | 12,6 | Small | Compact withdrawn | Dark green, no content | Thick filamented/ iridescent |

*(EtOH). Ethanol-fixed animals:* whole animal fixed before measurements in 96% Ethanol at field sampling. Examination data not applicable for gonads, foot and digestive gland when fixed.

**Supplementary Table S3.** Primers used in this study.
List of general and "Nematopsis clade" primers in this study (upper part) and schematic Figure of 18s operon indicating primer placement according to variable regions within the 18s rDNA operon (lower part).

| PRIMER/PROBE | SEQUENCE (5' – 3') | SPECIFICITY | REFERENCE |
|---|---|---|---|
| NonMet-F(18s-EUK581-F) | GTGCCAGCAGCCGCG | Eukaryote/Procaryote | Bower et al 2004 |
| NonMet-R(18s-EUK1134-R) | TTTAAGTTTCAGCCTTGCG | Non_Metazoan | Bower et al 2004 |
| IntPrimer F1 | GATTAAGCCATGCATGTCTAAG | Eukaryotes | Wakeman 2012 |
| Nem 1F_AA | CCCATGCTTGCATGGAGYKC | Nematopsis clade | This study |
| Nem2F_AA | CATTACTACACGGATACCTGTG | Nematopsis clade | This study |
| Nem3R_AA | TCCCGGTGCGAAATACAAA | Nematopsis clade | This study |
| Nem4R_AA | ATTTCTCCTGAGAACCCGGA | Nematopsis clade | This study |
| Nem5R_AA | CTGCAGGTTCACCTACAGGTA | Nematopsis clade | This study |
| 28sR1 | CGGTACTTGTTCGCTATCGG | Eukaryotes | Bass et al 2007 |
| M13F | GTAAAACGACGGCCAGT | Vector | |
| M13R | AGGAAACAGCTATGACCAT | Vector | |
| 360rev (ISH) | TGGACTGTTGCCAGTCCTTC | Lineage A (FPM) | This study |
| 1143rev (ISH) | TAGACGTATGATTGACGTGC | Lineage A (FPM) | This study |

The schematic figure shows the primers and direction (forward above and reverse below). Position 0-1700 of the operon are indicated with scales on top of the Figure. Hypervariable regions v1-v9 indicated with white boxes where approximate position v1 - v9 were taken from Hadziavdic et al. 2014.

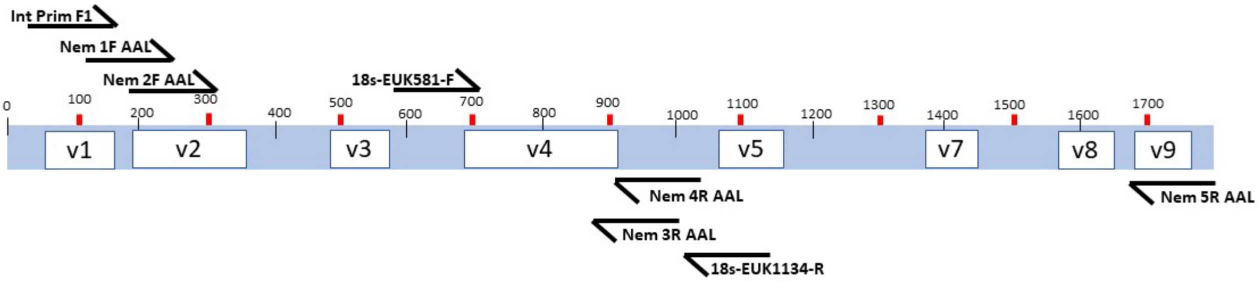

**Supplementary Table S4. Sequence generated in this study**

| Sequence name | phylogenetic assignament | locality | population type | collection year | sequencing technology | Accession number |
|---|---|---|---|---|---|---|
| FPM1_2019_MME_long_clone 13_17 | *Gregarinidae_sp* . lineage B | Sweden, Teakersälven, downstream river | MME | 2019 | Sanger | OR167184 |
| FPM5_2018_MME_long_clone12_19 | *Gregarinidae_sp.* lineage A1 | Sweden, Teakersälven, downstream river | MME | 2018 | Sanger | OR167028 |
| FPM4_2018_MME_long_clone11_19 | *Gregarinidae_sp.* lineage A1 | Sweden, Teakersälven, downstream river | MME | 2018 | Sanger | OR167022 |
| FPM4_2018_MME_long_clone11_20 | *Gregarinidae_sp.* lineage A1 | Sweden, Teakersälven, downstream river | MME | 2018 | Sanger | OR167021 |
| FPM4_2018_MME_long_clone11_5 | *Gregarinidae_sp.* lineage A1 | Sweden, Teakersälven, downstream river | MME | 2018 | Sanger | OR167024 |
| FPM4_2018_MME_long_clone11_9 | *Gregarinidae_sp.* lineage A1 | Sweden, Teakersälven, downstream river | MME | 2018 | Sanger | OR167023 |
| FPM5_2018_MME_nest_klonN5-5 | *Gregarinidae_sp.* lineage A1 | Sweden, Teakersälven, downstream river | MME | 2018 | Sanger | OR167030 |
| FPM2_2018_MME_nest_klon_N2_2 | *Gregarinidae_sp.* lineage A1 | Sweden, Teakersälven, downstream river | MME | 2018 | Sanger | OR167014 |
| FPM4_2018_MME_nest_klon59-4-7 | *Gregarinidae_sp.* lineage A1 | Sweden, Teakersälven, downstream river | MME | 2018 | Sanger | OR167027 |
| FPM9_2019_MME_nest_N16-6 | *Gregarinidae_sp.* lineage A1 | Sweden, Teakersälven, downstream river | MME | 2019 | Sanger | OR167194 |
| FPM5_2018_MME_nest_klonN5_6 | *Gregarinidae_sp.* lineage A1 | Sweden, Teakersälven, downstream river | MME | 2018 | Sanger | OR167031 |
| FPM4_2018_MME_nest_klonN4_1 | *Gregarinidae_sp.* lineage A1 | Sweden, Teakersälven, downstream river | MME | 2018 | Sanger | OR167026 |
| FPM5_2018_MME_nest_klonN5-4 | *Gregarinidae_sp.* lineage A1 | Sweden, Teakersälven, downstream river | MME | 2018 | Sanger | OR167029 |
| FPM3_2018_MME_Unonmet_klon_4_6 | *Gregarinidae_sp.* lineage A2 | Sweden, Teakersälven, downstream river | MME | 2018 | Sanger | OR167015 |
| FPM4_2020_MME_nest_N6 | *Gregarinidae_sp.* lineage A2 | Sweden, Teakersälven, downstream river | MME | 2020 | Sanger | OR167380 |
| FPM8_2019_MME_nest_N13_8 | *Gregarinidae_sp.* lineage A2 | Sweden, Teakersälven, downstream river | MME | 2019 | Sanger | OR167192 |
| FPM8_2019_MME_nest_N13_7 | *Gregarinidae_sp.* lineage A2 | Sweden, Teakersälven, downstream river | MME | 2019 | Sanger | OR167193 |
| FPM19_2020_MME_Upstream_nest_N21 | *Gregarinidae_sp.* lineage A2 | Sweden, Teakersälven, upstream river | MME | 2020 | Sanger | OR168122 |
| FPM15_2020_MME_Upstream_nest_N17 | *Gregarinidae_sp.* lineage A2 | Sweden, Teakersälven, upstream river | MME | 2020 | Sanger | OR168121 |
| FPM8_2020_MME_nest_N10 | *Gregarinidae_sp.* lineage A2 | Sweden, Teakersälven, downstream river | MME | 2020 | Sanger | OR167381 |
| FPM6_2019_MME_nest_N13_3 | *Gregarinidae_sp.* lineage A2 | Sweden, Teakersälven, downstream river | MME | 2019 | Sanger | OR167191 |
| FPM4_2019_MME_nest_N2_4 | *Gregarinidae_sp.* lineage A2 | Sweden, Teakersälven, downstream river | MME | 2019 | Sanger | OR167187 |
| FPM1_2019_MME_nest_N2-10 | *Gregarinidae_sp.* lineage A2 | Sweden, Teakersälven, downstream river | MME | 2019 | Sanger | OR167182 |
| FPM3_2018_MME_Unonmet_klon_4-3 | *Gregarinidae_sp.* lineage A2 | Sweden, Teakersälven, downstream river | MME | 2018 | Sanger | OR167020 |
| FPM6_2019_MME_nest_N13_4 | *Gregarinidae_sp.* lineage A2 | Sweden, Teakersälven, downstream river | MME | 2019 | Sanger | OR167190 |
| FPM5_2019_MME_nest_N2_5 | *Gregarinidae_sp.* lineage A2 | Sweden, Teakersälven, downstream river | MME | 2019 | Sanger | OR167189 |
| FPM4_2019_MME_nest_klonN4_4 | *Gregarinidae_sp.* lineage A2 | Sweden, Teakersälven, downstream river | MME | 2019 | Sanger | OR167188 |
| FPM4_2018_MME_nest_klonN4_2 | *Gregarinidae_sp.* lineage A2 | Sweden, Teakersälven, downstream river | MME | 2018 | Sanger | OR167025 |
| FPM3_2019_MME_nest_klonN15-1 | *Gregarinidae_sp.* lineage A2 | Sweden, Teakersälven, downstream river | MME | 2019 | Sanger | OR167186 |
| FPM2_2019_MME_nest_klon_N6-3 | *Gregarinidae_sp.* lineage A2 | Sweden, Teakersälven, downstream river | MME | 2019 | Sanger | OR167185 |
| FPM1_2019_MME_nest_klon_N1-4 | *Gregarinidae_sp.* lineage A2 | Sweden, Teakersälven, downstream river | MME | 2019 | Sanger | OR167183 |
| FPM3_2018_MME_long_clone10_19 | *Gregarinidae_sp.* lineage A2 | Sweden, Teakersälven, downstream river | MME | 2018 | Sanger | OR167016 |
| FPM3_2018_MME_long_clone10_8 | *Gregarinidae_sp.* lineage A2 | Sweden, Teakersälven, downstream river | MME | 2018 | Sanger | OR167017 |
| FPM3_2018_MME_long_clone10_3 | *Gregarinidae_sp.* lineage A2 | Sweden, Teakersälven, downstream river | MME | 2018 | Sanger | OR167019 |
| FPM3_2018_MME_long_clone10_6 | *Gregarinidae_sp.* lineage A2 | Sweden, Teakersälven, downstream river | MME | 2018 | Sanger | OR167018 |
| FPM5_2018_MME_nest_klon_N5-7 | *Gregarinidae_sp.* lineage A2 | Sweden, Teakersälven, downstream river | MME | 2018 | Sanger | OR167032 |
| FPM26_2020_REF_nest_N28 | *Gregarinidae_sp.* lineage A2 | Sweden, Stommabacken, creek | ref | 2020 | Sanger | OR168627 |
| FPM8_2018_REF_Unonmet_klon_9-6 | *Gregarinidae_sp.* lineage C | Sweden, Stommebacken, creek | ref | 2018 | Sanger | OR133340 |
| FPM8_2018_REF_Unonmet_klon_9-3 | *Gregarinidae_sp.* lineage C | Sweden, Stommebacken, creek | ref | 2018 | Sanger | OR133338 |
| FPM6_2018_REF_Unonmet_klon_7-5 | *Gregarinidae_sp.* lineage C | Sweden, Stommebacken, creek | ref | 2018 | Sanger | OR133333 |
| FPM7_2018_REF_Unonmet_klon_8-6 | *Gregarinidae_sp.* lineage C | Sweden, Stommebacken, creek | ref | 2018 | Sanger | OR133336 |
| FPM6_2018_REF_Unonmet_klon_7-2 | *Gregarinidae_sp.* lineage C | Sweden, Stommebacken, creek | ref | 2018 | Sanger | OR133332 |
| FPM7_2018_REF_Unonmet_klon_8-8 | *Gregarinidae_sp.* lineage C | Sweden, Stommebacken, creek | ref | 2018 | Sanger | OR133337 |
| FPM7_2018_REF_Unonmet_klon_8-1 | *Gregarinidae_sp.* lineage C | Sweden, Stommebacken, creek | ref | 2018 | Sanger | OR133334 |
| FPM8_2018_REF_Unonmet_klon_9-5 | *Gregarinidae_sp.* lineage C | Sweden, Stommebacken, creek | ref | 2018 | Sanger | OR133339 |
| FPM7_2018_REF_Unonmet_klon_8-2 | *Gregarinidae_sp.* lineage C | Sweden, Stommebacken, creek | ref | 2018 | Sanger | OR133335 |

**MME** = mass mortalities events; **ref** = reference (healthy) population

**Supplementary Table S5**. List of long-read *Nematopsis* sequences (eDNA) and their corresponding to environment and their origin from Sweden and Canada.

| LOCALITY | YEAR | DATE | REL. ABUN -DANCE | SEQENCE ID | COLLECTED BY | LATID. | LONGIT. | REFERENCE |
|---|---|---|---|---|---|---|---|---|
| **Sweden** | | | | | | | | |
| Lake Stortjärn | 2019 | 5 nov | 0,026 | c-4149_conseq _Otu094 | Swedish Infrastructure for Ecosystem Science | 64,24 | 19,76 | Jamy et al. (2022) |
| Lake Ersjön | 2019 | 17 oct | 0,0087 | c-8955_conseq _Otu0302 | Swedish Infrastructure for Ecosystem Science | 58,37 | 12,16 | Jamy et al. (2022) |
| Lake Erken | 2019 | 9 oct | 0,0018 | c-8783_conseq _Otu0423 | Swedish Infrastructure for Ecosystem Science | 59,83 | 18,63 | Jamy et al. (2022) |
| **Canada** | | | | | | | | |
| Permafrost thaw ponds | 2014 | 1 aug | 0,0019 | c-7557_conseq _Otu0783 | Sari Peura | 55,22 | -77,69 | Peura et al. (2020) |
| Permafrost thaw ponds | 2014 | 1 aug | 0,16 | c-7353_conseq _Otu0069 | Sari Peura | 55,22 | -77,69 | Peura et al. (2020) |
| Permafrost thaw ponds | 2014 | 1 aug | 0,038 | c-7549_conseq _Otu0168 | Sari Peura | 55,22 | -77,69 | Peura et al. (2020) |

**Supplementary Table S6. Accession numbers for the GenBank sequences used for the phylogenetic analyses.**

| GenBank number | Species | host | ENVIRONMENT |
|---|---|---|---|
| AB000912 | Tridacna hemolymph | clam | marine |
| AB252765.1 | anoxic sediment | eDNA | marine |
| AF009245 | Frenkelia glareoli | bankvole | terrestrial |
| AF013418.1 | Theileria parva | buffalo | terrestrial |
| AF022194.1 | Gymnodinium fuscum | free-living | freshwater |
| AF080611.1 | Lankesterella minima | amphibian | terrestrial |
| AF093489.1 | Cryptosporidium parvum | human | terrestrial |
| AF093499.1 | Cryptosporidium serpentis | snake | terrestrial |
| AF111187 | Cyclospora papionis | monkey | terrestrial |
| AF129882.1 | Gregarina niphandrodes | beetle | insect |
| AF129883.1 | Ophriocystis elektroscirrha | butterfly | insect |
| AF130361.1 | Hepatozoon catesbianae | american-bullfrog | terrestrial |
| AF286023.1 | Hematodinium sp | blue-crab | marine |
| AF298623 | Hyaloklossia lieberkuehni | amphibians | terrestrial |
| AF330214.1 | Microvorax tetrahymenae | freeliving | freshwater |
| AF399930.1 | Cytauxzoon felis | cat | terrestrial |
| AF457127 | Monocystis agilis | earthworm | terrestrial |
| AF494059.1 | Adelina bambarooniae | beetle | insect |
| AF497479.1 | Perkinsus marinus | oyster | marine |
| AJ271354.1 | Neospora caninum | mammal | terrestrial |
| AY078092 | Voromonas pontica | freeliving | marine |
| AY179975.1 | uncultured eukaryote eukaryote | eDNA | marine |
| AY179976.1 | uncultured eukaryote eukaryote | eDNA | marine |
| AY196706.1 | Lecudina polymorpha | annelids | marine |
| AY334568.1 | Mattesia geminata | ant | insect |
| AY490099.1 | undescribed eukaryote | flower-might | insect |
| AY603402.1 | Babesia bigemina | mite | terrestrial |
| AY821921.1 | Uncultured Eugregarinida | eDNA | freshwater |
| DQ093793.1 | Pterospora schizosoma | annelids | marine |
| DQ096836 | Adelina grylli | cricket | insect |
| DQ174731.1 | Chromera velia | cnidarians | marine |
| DQ176427.1 | Syncystis mirabilis | waterscorpion | insect |
| DQ273988 | Rhytidocystis polygordiae | polychaete | marine |
| DQ462457.1 | Ascogregarina culicis | mosquito | insect |
| DQ683562.1 | Selenidium serpulae | annelids | marine |
| EF024338.1 | uncultured eukaryote | eDNA | aspen rhizosphere elevated CO2 soil |
| EF024388.1 | uncultured eukaryote | eDNA | aspen rhizosphere elevated CO2 soil |
| EF024471.1 | uncultured eukaryote | eDNA | aspen rhizosphere elevated CO2 soil |
| EF100322.1 | uncultured eukaryote | eDNA | oxygen depleted sediment marine |
| EF492511.1 | Prorocentrum micans | freeliving | marine |
| EU162753.1 | Cryptosporidium fragile | amphibian | terrestrial |
| EU910605.1 | uncultured eukaryote | eDNA | sulfur rich hypoxic groundwater sediment freshwater |
| FJ459737.1 | Amoebogregarina nigra | grasshopper | insect |
| FJ459738.1 | Colepismatophila watsonae | silverfish | insect |
| FJ459739.1 | Geneiorhynchus manifestus | dragonfly | insect |
| FJ459741.1 | Gregarina blattarum | cockroach | insect |
| FJ459743.1 | Gregarina coronata | beetle | insect |
| FJ459746.1 | Gregarina kingi | grasshopper | insect |
| FJ459748.1 | Gregarina polymorpha | beetle | insect |
| FJ459750.1 | Hoplorhynchus acanthatholius | damselfly | insect |
| FJ459755.1 | Paraschneideria metamorphosa | fungus, gnats | terrestrial |
| FJ459756.1 | Prismatospora evansi | dragonfly | insect |
| FJ459761.1 | Stylocephalus giganteus | beetle | insect |
| FJ459762.1 | Xiphocephalus ellisi | beetle | insect |
| FJ459763.1 | Xiphocephalus triplogemmatus | beetle | insect |
| FJ832156.1 | Gregarinidae sp | peanutworm | marine |
| FJ832157.1 | Lecudina longissima | peanut-worm | marine |
| FJ832159.1 | Difficilina paranemertis | ribbonworms | marine |
| FJ832160.1 | Difficilina tubulani | ribbonworms | marine |
| FJ832161.1 | Selenidium orientale | peanutworm | marine |
| FJ832162.1 | Selenidium pisinnus | peanutworm | marine |
| FJ865354.1 | Psychodiella chagasi | sandfly | insect |

| GenBank number | Species | host | ENVIRONMENT |
|---|---|---|---|
| FN546182.1 | Apicystis bombi | bumbelbee | insect |
| GQ149767 | Rhytidocystis cyamus | polychaete | marine |
| GU320208.1 | Gregarina ctenocephali | catflea | insect |
| HQ876006.1 | Thiriotia pugettiae | graceful-kelp-crab | marine |
| HQ876007.1 | Heliospora caprellae | skeleton-shrimp | marine |
| HQ8911132 | Cephaloidophora cf | barnacle | marine sediment |
| JN857967.1 | Selenidium idanthyrsae | polychaete | marine |
| JX535336.1 | Polyplicarium citrusae | polychaete | marine |
| JX5353401 | Polyplicarium curvarae | polychaete | marine |
| JX5353441 | Polyplcarium lacrimae | polychaete | marine |
| JX5353481 | Polyplicarium translucidae | polychaete | marine |
| KC110869.1 | Selenidium sensimae | annelids | marine |
| KC110871.1 | Selenidium neosabellariae | polychaetes | marine |
| KT717636.1 | Nematopsis temporariae | common-frog | terrestrial |
| KT717637.1 | Nematopsis temporariae | common-frog | terrestrial |
| KT717638.1 | Nematopsis temporariae | common-frog | terrestrial |
| KT717639.1 | Nematopsis temporariae | common-frog | terrestrial |
| KT717640.1 | Nematopsis temporariae | common-frog | terrestrial |
| KT717641.1 | Nematopsis temporariae | common-frog | terrestrial |
| KT717642.1 | Nematopsis temporariae | common-frog | terrestrial |
| KT717643.1 | Nematopsis temporariae | European-tree-frog | terrestrial |
| KT717644.1 | Nematopsis temporariae | European-tree-frog | terrestrial |
| KT717645.1 | Nematopsis temporariae | European-tree-frog | terrestrial |
| KT717646.1 | Nematopsis temporariae | European-tree-frog | terrestrial |
| KT717647.1 | Nematopsis temporariae | European-tree-frog | terrestrial |
| KT717648.1 | Nematopsis temporariae | common-frog | terrestrial |
| KT717649.1 | Nematopsis temporariae | common-frog | terrestrial |
| KT717650.1 | Nematopsis temporariae | European-tree-frog | terrestrial |
| KT717651.1 | Nematopsis temporariae | European-tree-frog | terrestrial |
| KT717652.1 | Nematopsis temporariae | European-tree-frog | terrestrial |
| KT717653.1 | Nematopsis temporariae | European-tree-frog | terrestrial |
| KT717654.1 | Nematopsis temporariae | agile-frog | terrestrial |
| KT717655.1 | Nematopsis temporariae | agile-frog | terrestrial |
| KT717656.1 | Nematopsis temporariae | agile-frog | terrestrial |
| KT717657.1 | Nematopsis temporariae | agile-frog | terrestrial |
| KT717658.1 | Nematopsis temporariae | agile-frog | terrestrial |
| KT717659.1 | Nematopsis temporariae | agile-frog | terrestrial |
| KU664395.1 | Stylocephalus gigas | beetle | insect |
| L31843.1 | Pseudomonocystis lepidiota | undescribed species | insect |
| M64244.1 | Sarcocystis muris | birds | terrestrial |
| MH208617.1 | Colpodella sp | tick | insect |
| MK181532.1 | Quadruspinospora mexicana | grasshoper | insect |
| MK946169.1 | uncultured eukaryote | eDNA | soil |
| U40262.1 | Eimeria mitis | bird | terrestrial |
| X75453.1 | Toxoplasma gondii partial | mammal | terrestrial |

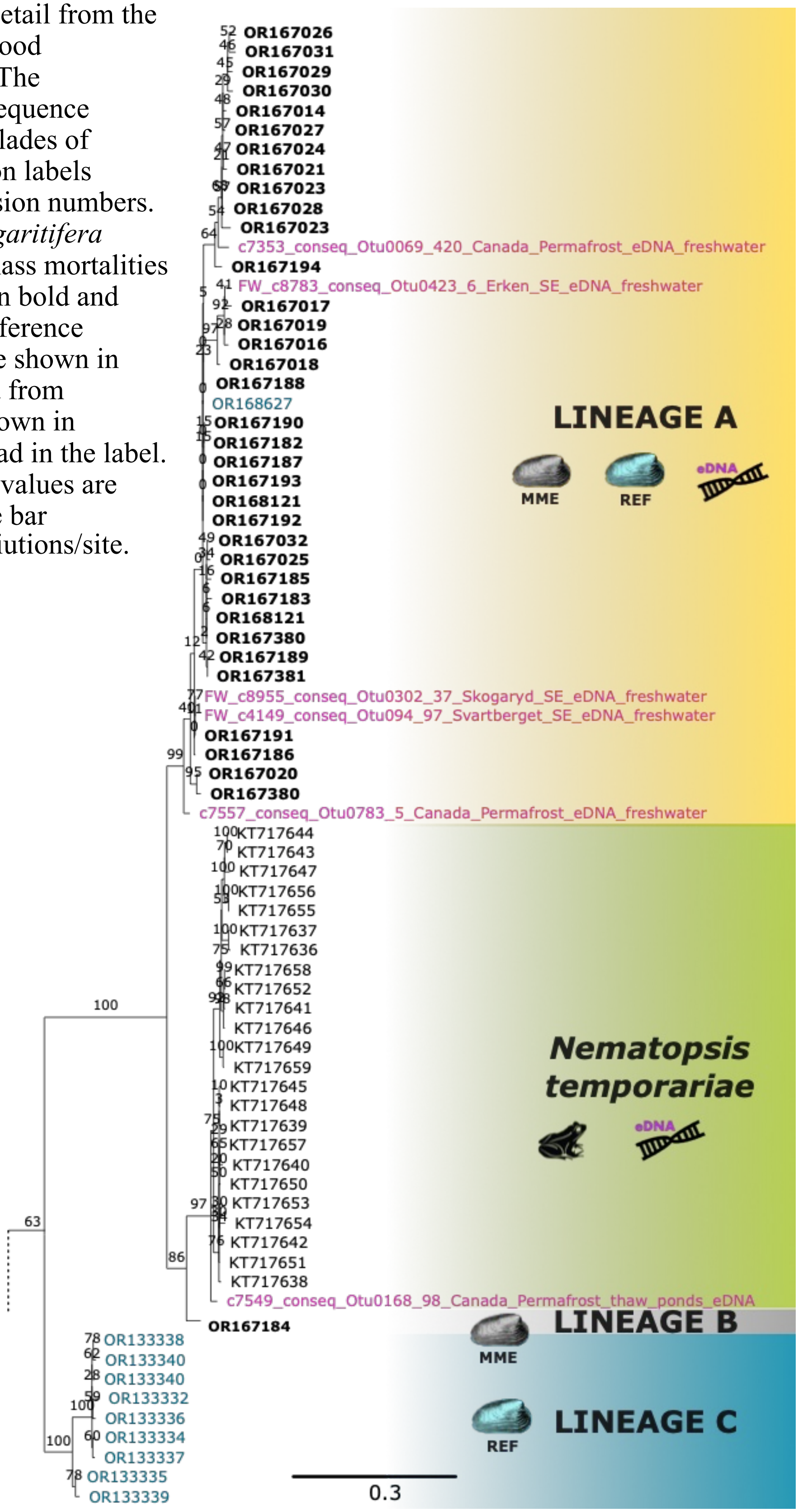


**Supplementary figure S1**: Detail from the 18S rDNA Maximum Likelihood phylogenetic tree (Figure 1). The phylogeny is displaying the sequence diversity of the uncollapsed clades of Lineage A, B and C. The taxon labels represent the GenBank accession numbers. The sequences of the *M. margaritifera* populations that undergone mass mortalities events (MME) are in shown in bold and black while the ones of the reference healthy populations (REF) are shown in blue. The sequences retrieved from environmental datatset are shown in purple. Their origin can be read in the label. The non-parametric boostrap values are shown on branches. The scale bar represents the number substitiutions/site.

**Supplementary figure S2**: Macroscopical observations of freshwater pearl mussels collected from ongoing mortality in Teåkersälven, with focus on gonad development. (A). Mussels collected 2018 in September, showing signs of thickening of mantel (yellow arrow). (B) The only mussel (M2-20) within MME population 2020 June, where macroscopic signs of gonad development (yellow arrowhead) were visual. (C) Mussel without signs of gonad development (M18-20).

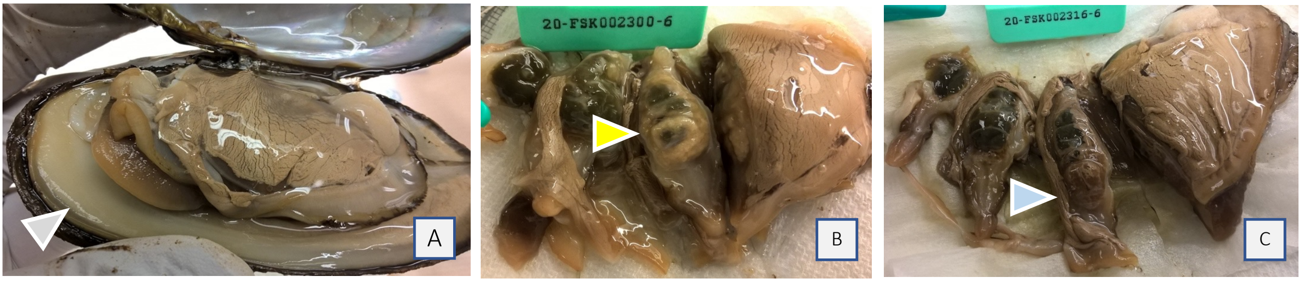

**Supplementary Figure S3**: Histology, with observations of lesions and degenerative changes within the gill, mantel, heart and digestive glands. (A) Hemocytes in mantel phagocytizing weakly eosinophilic microcells, resembling gymnospores or spheric zoites, dislocating the host cell nuclei, indicated with white arrowhead. (B) Aggregating hypertrophied hemocytes within mantel, some disruptered, releasing infiltrating cells. (C) Macrophagic cells within heart auricle, filled with granular content resembling pigmented invading spherical zoites or gymnospores. (D) Histozoic cyst with folded vermozoite cells within connective tissue enclosing the hemolymph dorsal compartment. (E) Release of histozoic cyst from the mantel. (F) Disruption and release of epithelial cell within digestive gland. (G) Ruptered areas in the digestive gland epithelia layer, leaving empty spaces or caverns. (H) Formation of sporoblast and oocyst within digestive gland epithelial cells, some with openings towards lumen of the digestive gland. Section 4μm in thickness, stained with hematoxylin eosin, cellsizes indicated with scalebar, all in size 10μm.

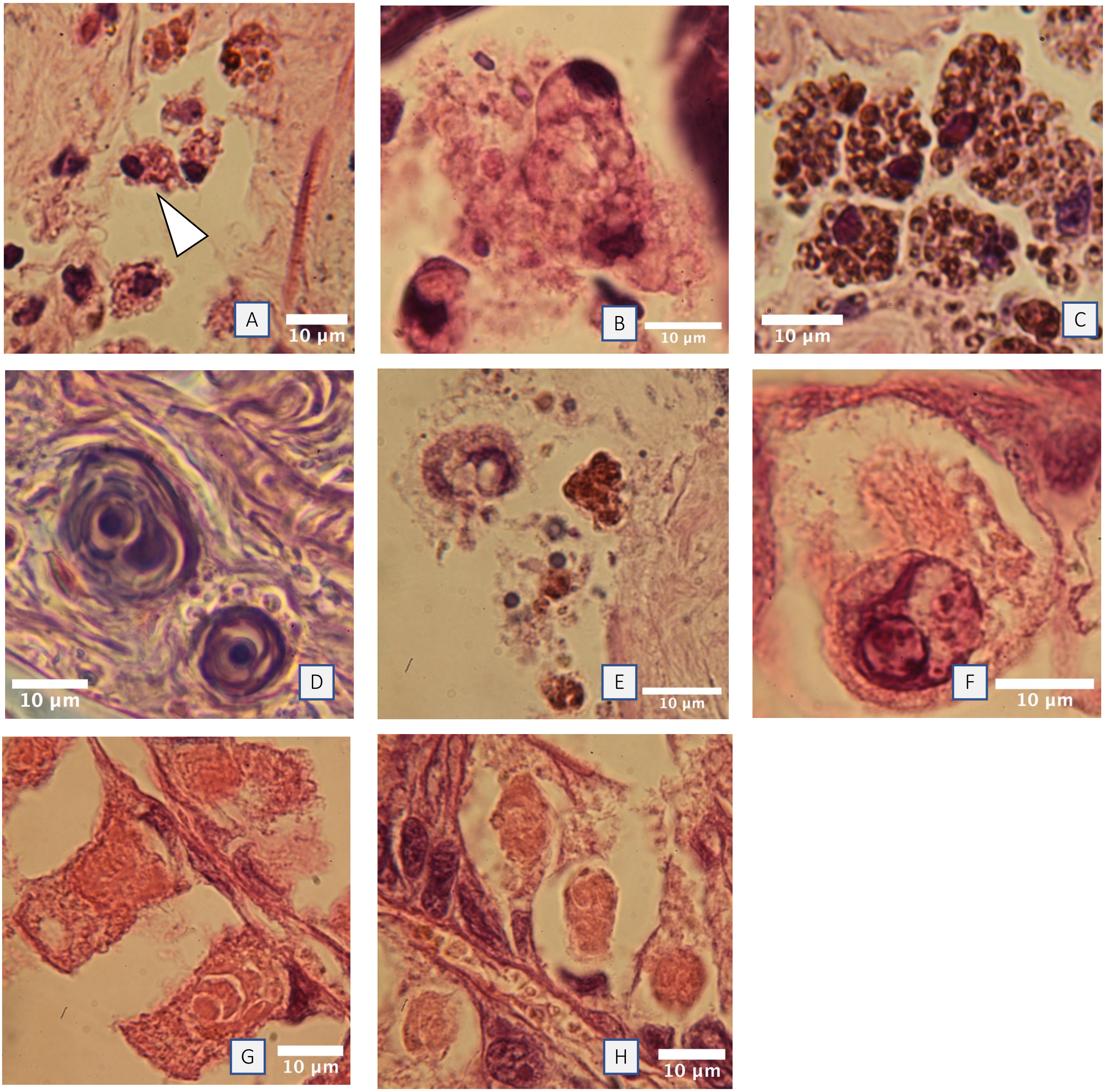

**Supplementary Figure S4**: Cellmorphology of infiltrating gregarine cells within the gill, mantel and digestive glands. (A) Singel spherical zoite within a thin transparent vacuole, embedded within hemolymph rich area of the mantel. (B) Enclosed weakly stained eosinophilic zoites within host-cell or gymnospore. (C-E) multiple spherical zoites enclosed within hemolymph droplets, some also released in the dorsal hemolymph compartment. (F) Histozoic basophilic zoites cysts enclosed within connective tissue of the mantel.
(G) Three histozoic basophilic cysts within the connective tissue of gills, imaged by phase contrast, revealing single or multiple folded zoites within. (H) Multiple free zoites transported down into the digestive glands via duct channels. (I) Two differently stained zoites within digestive glands in syzygy like, head to head, formation in the digestive glands epithelium, phase contrast image. (J) Release of oocyst or sporocyst from parasitophorous vacuole within digestive gland of digestive gland, phase contrast. (K) Singel oocyst or sporoblasts enclosed within parasitophorous vacuole in the digestive gland epithelium. Section 4µm in thickness, stained with hematoxylin eosin, cell-sizes indicated with scalebar, 5µm (A, E), 10µm (B-D, F, G, J, K) and 20µm (H, I).

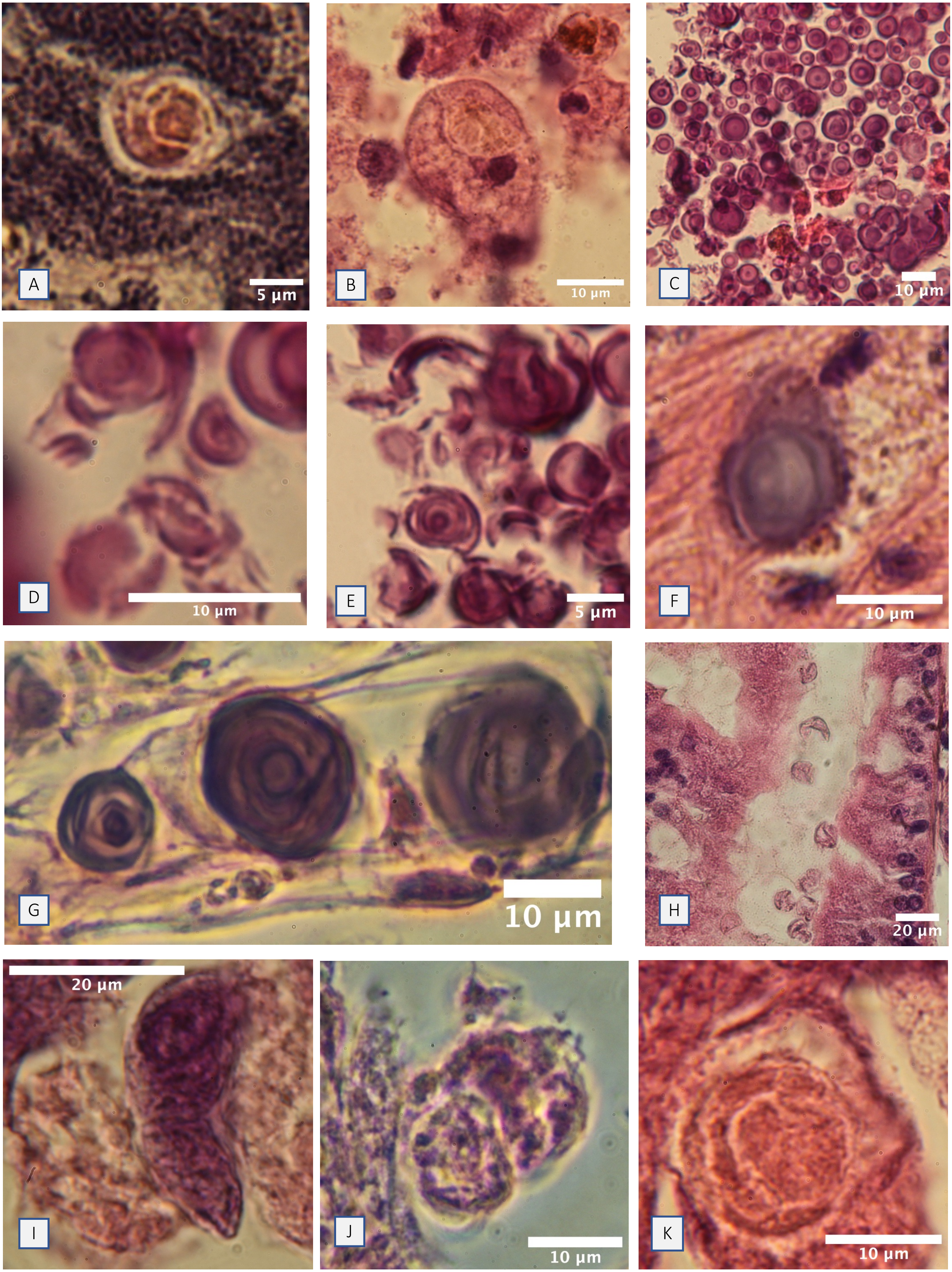
A
5 µm
B
10 µm
C
10 µm
D
10 µm
E
5 µm
F
10 µm
G
10 µm
H
20 µm
I
20 µm
J
10 µm
K
10 µm
